# Cell-type-specific sustained value representations in the claustrum

**DOI:** 10.64898/2026.03.10.710905

**Authors:** Ahmad B. Taha, Seong Yeol An, Su-Jeong Kim, Rowhan Daly, Jeremiah Y. Cohen, Solange P. Brown

**Affiliations:** Solomon H. Snyder Department of Neuroscience, Johns Hopkins University School of Medicine, Baltimore, MD 21205, USA; Kavli Neuroscience Discovery Institute, Johns Hopkins University School of Medicine, Baltimore, MD 21205, USA; Allen Institute for Neural Dynamics, Seattle, WA 98109, USA

## Abstract

Flexible decision-making relies on interactions between frontal cortex and subcortical structures. The claustrum, a subcortical nucleus highly interconnected with frontal cortex, influences cortical activity and has been implicated in cognitive functions. Recording from claustrum neurons as mice performed a reinforcement learning task, we found that the activity of almost half of recorded neurons scaled with reward rate and predicted trial-by-trial adjustments in reaction time and choice switching. Individual neurons sustained this activity over seconds between trials. Our recordings identified two electrophysiologically distinct populations. One was excited during task execution and bidirectionally scaled its activity with reward rate. The other was suppressed during task execution, scaled activity inversely with reward rate and projected to frontal cortex, indicating that claustrocortical outputs produce graded increases in activity with decreasing reward rate. Our results identify the claustrum as a subcortical locus for stable value representations and integrate it into neuronal circuits for value-based decision-making.

## Introduction

Flexible behavior depends on neocortex (Gold and Shadlen, 2007; Wang, 2008; Lee et al., 2012; Hanks and Summerfield, 2017; Guo et al., 2017a; Uddin, 2021). Frontal cortex is central to this flexibility, with its activity tracking information about recent experience that animals use to inform future decisions (Miller and Cohen, 2001; Barraclough et al., 2004; Kennerley et al., 2006; Sul et al., 2010; Tsutsui et al., 2016; Bari et al., 2019; Hattori et al., 2023). This cortical activity emerges from interactions within distributed cortical networks and with subcortical structures, such as the thalamus (Minamimoto et al., 2005; Browning et al., 2015; Rikhye et al., 2018; Guo et al., 2017b; Yang et al., 2022). The claustrum is another subcortical nucleus that is highly interconnected with frontal cortex (Clascá et al., 1992; Smith and Alloway, 2010; Torgerson et al., 2015; Atlan et al., 2017; Wang et al., 2017; White et al., 2017; Zingg et al., 2018; Chia et al., 2020; Marriott et al., 2021; Wang et al., 2023), and several lines of evidence suggest that it may be involved in dynamic decision making. First, modulating the activity of claustrocortical projections influences the activity of frontal cortex (Jackson et al., 2018; Narikiyo et al., 2020; Fodoulian et al., 2020; McBride et al., 2023; Atilgan et al., 2025; De La Torre-Martínez et al., 2025; Peretz-Rivlin et al., 2026) and has implicated claustrocortical neurons in higher cognitive functions in behaving animals, including attention (Atlan et al., 2018; White et al., 2018; Terem et al., 2020; Fodoulian et al., 2020), impulsivity (Liu et al., 2019; Atlan et al., 2024) and learning (Grasby and Talk, 2013; Reus-García et al., 2021; Atilgan et al., 2025). Second, cortical neurons in the dorsomedial frontal cortex projecting to the claustrum encode expected value and economic risk (Majumdar et al., 2025). Third, claustrum activity may persist for many seconds during behavior. For example, individual claustrum neurons exhibit sustained activity during intertrial intervals (ITIs), predictive of whether an animal will lick left or right for water rewards associated with sensory cues (Chevée et al., 2022). Some studies have also implicated sustained population activity across claustrum neurons in behaviors that depend on working memory (Han et al., 2024; Bhattacharjee et al., 2024). These observations raise key questions: does the claustrum represent sustained value signals during dynamic decision making, and if so, what information does it convey to frontal cortex during flexible behavior?

To test whether claustrum activity represents value signals, we recorded the electrophysiological activity of claustrum neurons in mice performing a dynamic foraging task that required adapting behavior as reward probabilities associated with alternative actions changed. Claustrum neuron firing rates represented licking, choice and outcome during task execution. During the ITIs, after licking had ceased, approximately 40% of neurons exhibited activity that correlated with a continuously updated estimate of total value in the environment, reflecting reward rate. The activity of individual neurons stably represented these total value estimates over many seconds during ITIs and predicted trial-by-trial adjustments in behavior. We further identified two electrophysiological types of claustrum neurons and compared their response properties. One type was suppressed during active task performance, and preferentially scaled its activity inversely with total value during the ITIs. We show that this type includes neurons projecting to medial frontal cortex. The other type, in contrast, was generally excited during task performance, had comparable fractions of neurons with activity positively or negatively correlated with total value and did not project to frontal cortex. These results identify the claustrum as a subcortical locus of persistent value signals and suggest that claustrocortical neurons, which exhibited graded, monotonic increases in activity with decreasing total value, convey continuously updated estimates of reward availability to medial frontal cortex. Together, our data suggest that the claustrum is integrated into the neural circuits that support flexible decision making.

## Results

### Claustrum neurons represent choice and outcome during dynamic foraging

To test whether claustrum activity represents value signals during flexible decision making, we first trained head-restrained mice on a dynamic foraging task in which the reward contingencies of two water ports fluctuated independently over time (Fig. 1a) (Grossman et al., 2022; Su and Cohen, 2022). Following an auditory go cue, mice freely chose to lick the left or right port, and water rewards were delivered probabilistically (*P* (*R*) ∈ {0.1, 0.5, 0.9}) after a fixed 300 ms delay. Choices were followed by variable ITIs of up to 21 seconds. Reward probabilities for each port changed pseudorandomly and independently every 20–35 trials. In 10% of trials, an auditory no-go cue was presented and no reward was delivered. Well-trained mice responded to the go cue on 98% of trials (IQR = 95–99% per session, *n* = 119 sessions, *N* = 9 mice) and preferentially selected the higher-probability option with minimal side bias (Fig. S1a,b; *n* = 119 sessions, *N* = 9 mice; Fraction of optimal choices = 0.66±0.021), achieving rewards on 0.56±0.018 of trials. Mice adapted their behavior to track the underlying reward contingencies (Fig. 1b), with choice allocation closely following reward availability (Fig. 1c; *R*^2^ = 0.83; slope = 0.78 ± 0.03). Following changes in reward probability, choices changed over several trials, with faster and larger shifts observed under greater reward asymmetries (Fig. S1c) (Grossman et al., 2022). Logistic regression analyses further revealed that upcoming behavioral choices depended on multiple past outcomes, with recent outcomes exerting a stronger influence than more distant outcomes (Fig. 1d, Fig. S1d), indicating that mice learned continuously from recent experience and adjusted their decisions to match the underlying reward environment.

**Figure 1.**
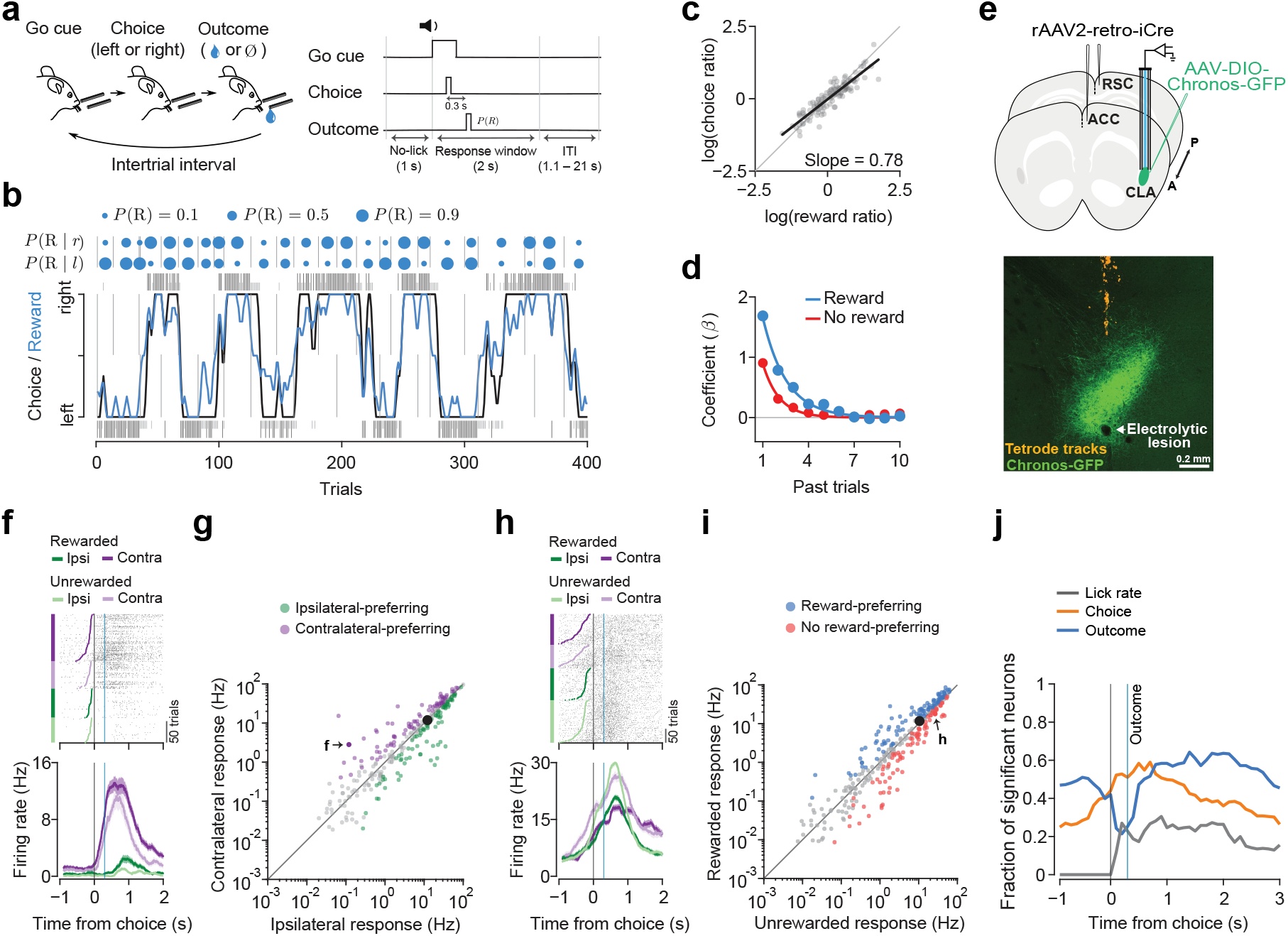
Claustrum neurons represent choice and outcome during a dynamic foraging task. (**a**) Schematic of the dynamic foraging task. Following an auditory cue, head-fixed mice choose between left or right lick ports. A probabilistic water reward is delivered after a 300 ms delay. Reward probabilities at each port change independently every 20–35 trials throughout the behavioral session. (**b**) Example behavioral session from a well-trained mouse. Blue circles denote reward probabilities, with dashed lines marking probability changes. The black trace shows smoothed choices, and the blue trace shows smoothed rewards. Ticks (Tall black: rewarded; Short gray: unrewarded) indicate right (above) and left (below) choices. (**c**) Scatter plot of the log-ratio of rewards obtained and choices made across sessions (*n* = 119 sessions, *N* = 9 mice). Gray circles represent individual sessions. The black line shows the linear regression fit (*R*^2^ = 0.83, *slope* = 0.78 ± 0.03), and the gray diagonal indicates perfect matching. (**d**) Logistic regression coefficients predicting upcoming choice from past outcome history (Blue: coefficients for past rewarded trials; Red: coefficients for past unrewarded trials). Solid curves show exponential fits; error bars (95% CI) are obscured by the symbols. (**e**) Top, experimental configuration. C57BL/6J mice were injected with rAAV2-retro-iCre in ACC and RSC and AAV2-FLEX-Chronos-GFP in the left claustrum, and implanted with an optic fiber and tetrodes coated with a fluorescent dye. Bottom, tetrode tracks (orange) through the claustrum (green, Chronos-GFP). The arrow indicates an electrolytic lesion. Scale bar: 0.2 mm. (**f**) Raster plots and PSTHs of an example neuron showing stronger responses for contralateral choices (Purple: contralateral; Green: ipsilateral), aligned to choice onset. Blue line: outcome delivery. Colored dots in the raster indicate cue onset. (**g**) Scatter plot of claustrum neuron firing rates during ipsilateral versus contralateral choices (*n* = 341 neurons, *N* = 9 mice), computed from −100 to 100 ms around the first lick (Green: ipsilateral-preferring; Purple: contralateral-preferring neurons). Black dot represents the population mean (Ipsilateral: 12.2 ± 0.99 Hz, Contralateral: 11.9 ± 0.97 Hz; *p* = 0.25, Wilcoxon signed-rank test). Error bars (SEM) are obscured by the symbol. Dot denoted as f shows the example neuron in (f). (**h**) Raster plots and PSTHs of an example neuron showing stronger responses on unrewarded trials (Dark: rewarded; Light: unrewarded), aligned to choice onset. Blue line: outcome delivery. Colored dots in the raster indicate cue onset. (**i**) Scatter plot of claustrum neuron firing rates for rewarded versus unrewarded trials (*n* = 341 neurons, *N* = 9 mice), computed during the first 500 ms after outcome delivery (Blue: reward-preferring; Red, no reward-preferring neurons). Black dot represents the population mean (Rewarded: 11.7 ± 0.93 Hz; Unrewarded: 10.5 ± 0.80 Hz; *p* = 0.058, Wilcoxon signed-rank test). Error bars (SEM) are obscured by the symbol. Dot denoted as h shows the example neuron in (h). (**j**) Fraction of neurons with significant regression coefficients from linear models relating firing rates to lick rate, choice, and outcome, computed in 100-ms bins aligned to choice (*n* = 341 neurons, *N* = 9 mice). Outcome delivery is indicated by the blue vertical line.

We recorded action potentials from 341 neurons in the anterior claustrum of nine mice performing the dynamic foraging task using tetrode bundles tethered to an optic fiber (Fig. S1e–g), verifying tetrode placement within the claustrum using three complementary approaches (Fig. 1e, Fig. S1h,i; see Methods) (Chevée et al., 2022). We first analyzed neuronal activity during task execution, when mice made choices and outcomes were revealed. The activity of the vast majority of neurons was significantly modulated on go trials, when mice responded following the auditory cue (Fig. S1j; 89%, *n* = 305 of 341 neurons, Wilcoxon signed-rank test). Few neurons responded during no-go trials in the absence of licking (Fig. S1k,l; 8%, *n* = 15 of 180 neurons; Wilcoxon signed-rank test), consistent with prior studies reporting few sensory-only responses in anterior claustrum (Ollerenshaw et al., 2021; Chevée et al., 2022). Almost half of recorded neurons responded significantly more to one lick direction than the other (Fig. 1f,g; 48%, *n* = 164 of 341 neurons, Wilcoxon rank-sum test), with no significant population bias in preference for ipsilateral versus contralateral lick direction (Ipsilateral: 12.2 ± 0.99 Hz, *n* = 88 of 341 neurons; Contralateral: 11.9 ± 0.97 Hz, *n* = 76 of 341 neurons; *p* = 0.25, Wilcoxon signed-rank test), consistent with prior work using a sensory selection task (Chevée et al., 2022). The activity of most neurons also differed significantly between rewarded and unrewarded trials (Fig. 1h,i; 58%, *n* = 197 of 341 neurons, Wilcoxon rank-sum test), with no significant population bias for either outcome (Rewarded: 11.7 ± 0.93 Hz, *n* = 104 of 341 neurons; Unrewarded: 10.5 ± 0.80 Hz, *n* = 93 of 341 neurons; *p* = 0.058, Wilcoxon signed-rank test). Using linear regression, we found that most neurons had significant coefficients for lick rate, choice, outcome or their combinations (Fig. S1m,n). On average, mice exhibited more licks on rewarded than unrewarded trials (Fig. S1m; Rewarded: 9.7 ± 0.1 licks, Unrewarded: 4.1 ± 0.06 licks, *n* = 119 sessions; paired *t*- test, *p* = 1.18 ×10^−83^), raising the possibility that differences in neural activity correlated with outcome reflect differences in motor behavior associated with reward delivery. To account for this possibility, we decoded outcome using linear support vector machines (SVMs) from either raw firing rates or firing rates residualized to remove variance explained by licking and choice. Outcome was significantly decoded for 17% of neurons after residualization (*n* = 57 of 341 neurons), consistent with outcome-related signals not explained by licking and choice (Fig. S1o; see Methods).

To determine how neural representations of lick rate, choice, and outcome evolve during task execution, we fit time-resolved linear models that predicted neuronal firing rates from these task variables (Fig. 1j). The activity of approximately 20% of neurons correlated with lick rate shortly after the onset of licking. Over 50% of neurons showed significant choice coefficients following choice onset and approximately 20% did so after licking had ceased. Following reward delivery, more than 50% of neurons exhibited significant coefficients for outcome, with this fraction remaining above 40% beyond the period during which mice licked to receive feedback about the trial outcome. Together, these results indicate that choice and outcome representations evolve during task execution and persist after licking has ceased, extending into the ITI (Fig. S1p).

### Claustrum neuron activity during the intertrial intervals represents history signals

As we hypothesize that the claustrum represents information about prior experience between trials, we next asked whether claustrum neuron activity during the ITIs reflects prior choice and outcome. Applying a similar linear model as in Fig. 1j to the 1-s lick-free window preceding the go cue, a period when the lick rate is zero and the timing of the upcoming trial is not predictable, we found that more than half of claustrum neurons had significant coefficients for prior choice, prior outcome or both (Fig. 2a). When applying a regression model to the average firing rates for this same 1-s window, the activity of most recorded neurons was significantly correlated with the most recent outcome (Fig. 2b; 38%, *n* = 128 of 341 neurons) or with the combination of the most recent outcome and choice (25%, *n* = 86 of 341 neurons), while fewer neurons showed significant correlations with the most recent choice alone (12%, *n* = 41 of 341 neurons). These correlations were not simply restricted to the immediately preceding trial. When we fit the activity of each neuron during the 1-s lick-free window with a single regression model containing lagged choices, outcomes, and choice-outcome interaction terms spanning the past five trials, we found that 28% of neurons had significant coefficients for the most recent choice or choice-outcome interaction while only ∼5% had significant coefficients for choice for more than one trial back (Fig. S2a). In contrast, a significantly larger population of neurons had significant coefficients for outcome across trial lags: 44% of neurons had a significant coefficient for outcome at *t* − 1, 23% at *t* − 2, and 10% at *t* − 5 (exact permutation test, *p* = 0.008; post hoc sign-flip tests: *p* = 0.031). To quantify the span of trial history that best explains the activity of individual neurons, we used a nested model comparison, sequentially adding choice, outcome, and interaction terms from additional trials in the past (Fig. S2b; see Methods). Relative to models containing only *t* − 1 terms, adding *t* − 2 terms significantly improved fit for 52% of neurons (Fig. S2c; 131 of 254 neurons), with some neurons showing further improvement when terms from more distant trials were included (*t* − 3: 30%, 77 of 254; *t* − 4: 14%, 35 of 254; *t* − 5: 7%, 19 of 254).

**Figure 2.**
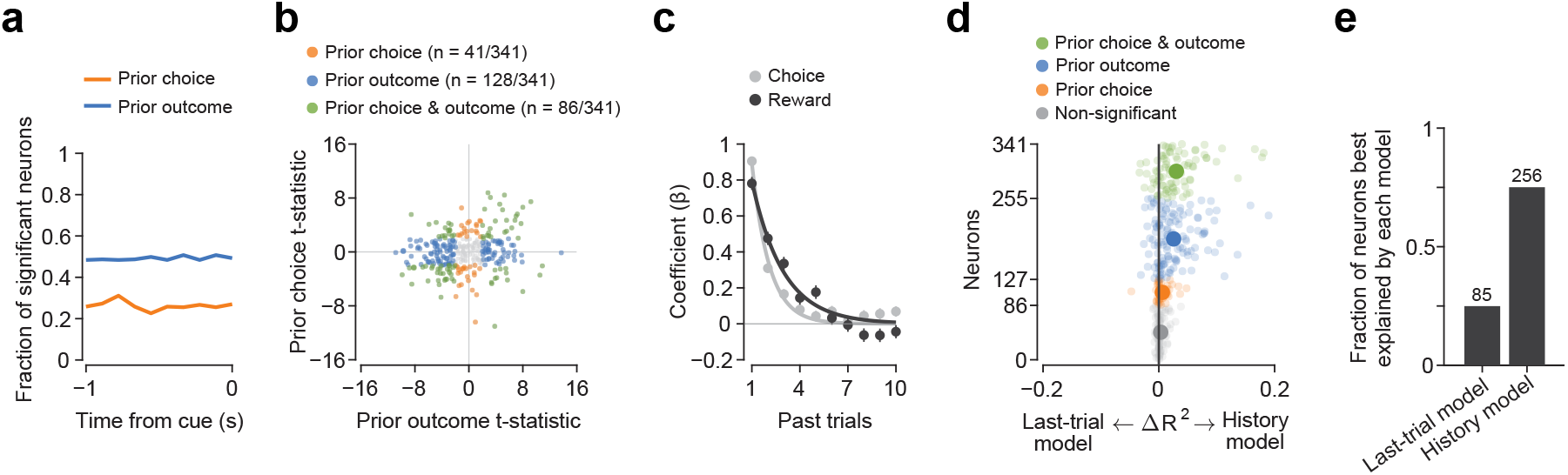
Claustrum neuron activity during the intertrial interval (ITI) represents history signals. (**a**) Fraction of neurons with significant regression coefficients from linear models relating firing rates during the 1-s lick-free pre-cue window to prior choice (orange) and prior outcome (blue), computed in 100-ms bins (*n* = 341 neurons). (**b**) Scatter plot of *t*-statistics from a single regression model fit to firing rates averaged across the 1-s pre-cue window for all 341 neurons. Neurons with significant coefficients for prior choice (orange, *n* = 41*/*341 neurons), outcome (blue, *n* = 128*/*341 neurons), or both (green, *n* = 86*/*341 neurons) are indicated. (**c**) Logistic regression coefficients predicting upcoming choice from past choice and reward history. Gray: coefficients for past choices. Black: coefficients for past rewards. Solid curves show exponential fits. Error bars: 95% CI. (**d**) Plot of the difference in variance explained (Δ*R*^2^) for firing rates in the 1-s pre-cue window between a model using the immediately preceding trial choice and outcome (Last-trial model) and a model using choice and reward history (History model), where history predictors were computed as a sliding exponential average over the past 10 trials using coefficient fits from (c). Positive Δ*R*^2^ values indicate neurons with activity better explained by the History model. Neurons are color-coded as in (b). (**e**) Fraction of neurons best explained by each model based on Akaike Information Criterion (AIC).

The presence of trial history representations in claustrum activity during ITIs raises a key question: do these neural signals align with the behavioral variables that guide choice? The mice’s decisions showed gradually decaying influences of reward history as well as choice history (Fig. 2c). To test whether claustrum neuron activity during the ITIs reflects these behaviorally relevant variables, we compared a model that includes only the choice and outcome of the immediately preceding trial (Last-trial model) with a model in which these predictors were replaced by exponentially weighted choice and reward history terms derived from fits to the behavioral data (History model; see Methods). We found that activity in 75% of neurons was better explained by the History model, despite the two models having the same number of parameters (Fig. 2d,e; Last-trial model: 85 of 341 neurons; History model: 256 of 341 neurons; Akaike Information Criterion). This improvement was driven primarily by reward history, which explained significantly more unique variance than choice history (Fig. S2d; Wilcoxon signed-rank test, *p <* 0.001). Together, these results indicate that claustrum neurons represent history signals which guide upcoming choices (Fig. 2c).

### Sustained claustrum activity during the intertrial intervals correlates with total value

In reinforcement-learning (RL) frameworks, histories of rewarded and unrewarded actions are modeled as action value estimates that evolve gradually over trials to inform upcoming decisions (Rescorla and Wagner, 1972; Bertsekas and Tsitsiklis, 1996; Sutton and Barto, 1998). Given that claustrum neurons integrate information over multiple past trials, we hypothesized that claustrum neuron activity may represent action values (*Q*_*l*_, *Q*_*r*_: corresponding to the left and right actions) and their combinations: total value (Σ*Q* = *Q*_*l*_ + *Q*_*r*_) and relative value (Δ*Q* = *Q*_*l*_ − *Q*_*r*_). We therefore fit a *Q*-learning model to behavior (Fig. 3a; Fig. S3a). Using firing rates in the 1 s prior to the go cue, we regressed neural activity onto *Q*_*l*_, *Q*_*r*_, and choice, and classified value signals based on the relative magnitude of regression coefficients in value space (Seo et al., 2009; Wang et al., 2013). Claustrum neurons with significant coefficients primarily aligned along the Σ*Q* axis (Fig. 3b), indicating that the activity of almost half of the recorded claustrum neurons was significantly correlated with Σ*Q* (Fig. 3c; 43%, *n* = 147 of 341 neurons), comprising two subsets with activity scaling positively or negatively with Σ*Q* (Fig. S3b; Σ*Q*^+^: *n* = 60*/*341 neurons; Σ*Q*^−^: *n* = 87*/*341 neurons). Individual neurons exhibited monotonic relationships between firing rate and Σ*Q* (Fig. 3d,e), and their trial-by-trial activity closely tracked Σ*Q* (Fig. 3f; *r* = −0.50). Claustrum Σ*Q* signals were not action-specific, as the relationship between firing rates and Σ*Q* was similar whether mice made.ipsilateral or contralateral choices in the upcoming (Fig. S3c, left; paired *t*-test, *t* = −1.5, *p* = 0.16) or past trials (Fig. S3c, right; paired *t*-test, *t* = −0.6, *p* = 0.56).

**Figure 3.**
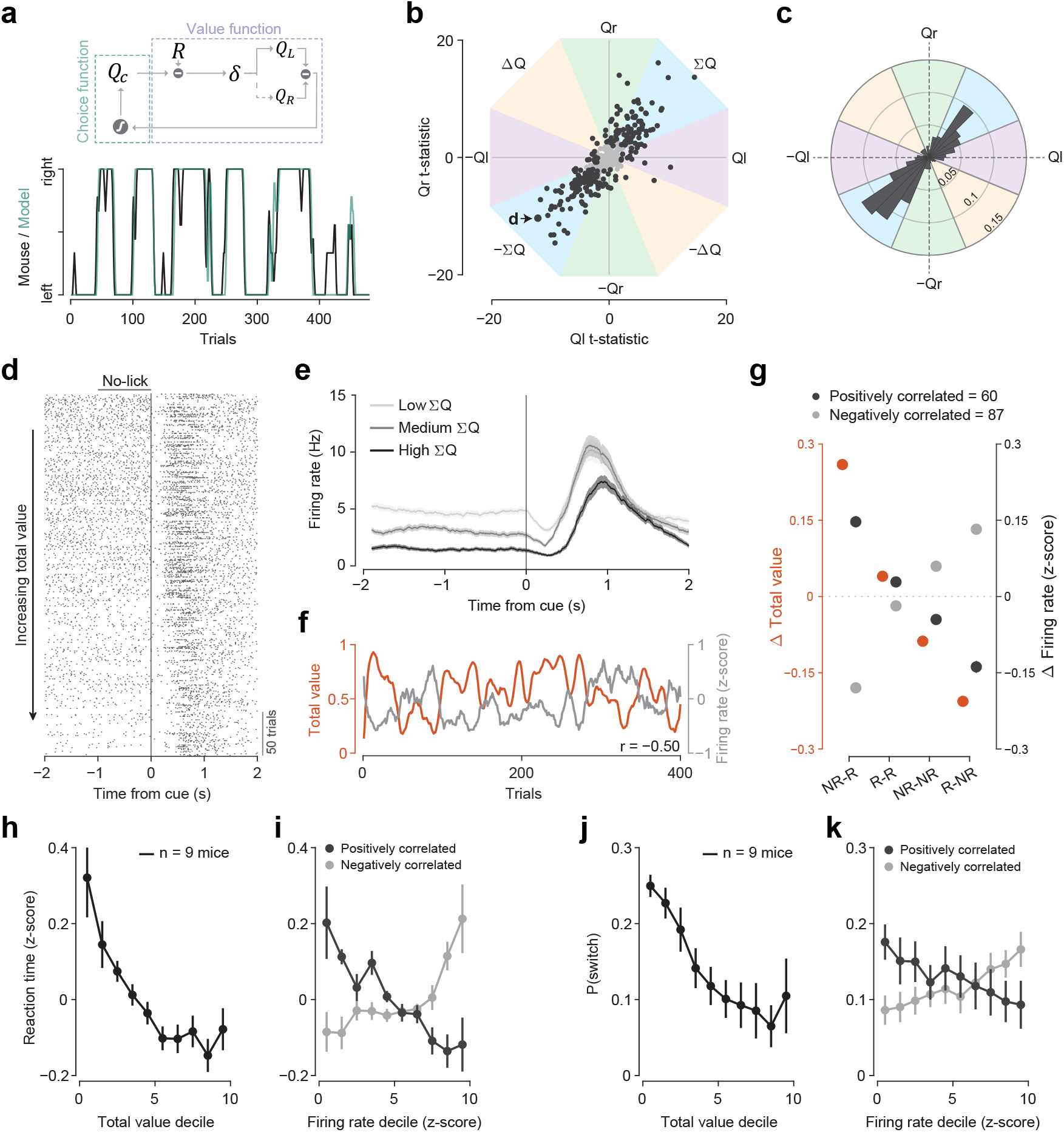
Claustrum neuron activity during the intertrial intervals (ITIs) scales with total value and predicts reaction time and probability of switching. (**a**) Top, schematic of the *Q*-learning model. The value function estimates action values for the left and right options (*Q*_*l*_, *Q*_*r*_) and their arithmetic combinations, total (Σ*Q*) and relative (Δ*Q*) value, to inform the next choice (*Q*_*c*_). Reward prediction errors (*δ*), scaled by learning rate (*α*), update action values. Bottom, *Q*-learning fit (green) to example mouse behavior (black). (**b**) Scatter plot of *t*-statistics from a linear model correlating firing rates to *Q*_*l*_, *Q*_*r*_, and choice projected into value space (*n* = 341 neurons). Colored sectors indicate value-coding subspaces (*Q*_*l*_, *Q*_*r*_, Σ*Q*, Δ*Q*). Black points denote neurons with significant coefficients. The point labeled d denotes the example neuron shown in (d–f). (**c**) Polar histogram of all neurons across value space, showing an over-representation along the Σ*Q* axis (*n* = 341 neurons, *V* -test on folded angles, *p <* 0.001). (**d**) Example Σ*Q* neuron. Cue-aligned raster plot of trials sorted by Σ*Q* shows decreasing pre-cue firing with increasing Σ*Q*. (**e**) PSTHs for the neuron in (d), split into terciles of low, medium, and high Σ*Q* trials. (**f**) Trial-by-trial comparison showing a negative correlation (*r* = −0.50) between model-derived Σ*Q* and neuronal firing rate for the example neuron in (d). (**g**) Comparison of model-predicted changes in Σ*Q* (orange) and measured changes in *z*-scored neuronal firing rates across the four possible outcome transitions between two consecutive trials (NR→R, R→R, NR→NR, R→NR; NR = no reward, R = reward). Means ± SEM are shown for neurons whose firing rates are significantly positively correlated (Σ*Q*^*+*^: dark gray; *n* = 60*/*341 neurons) or inversely correlated (Σ*Q*^−^: light gray, *n* = 87*/*341 neurons) with Σ*Q*. (**h**) Reaction time decreases monotonically with increasing Σ*Q* (*r* = −0.93, *p* = 1.2 ×10^−4^). Points show mean ± SEM (*n* = 9 mice). (**i**) Pre-cue firing rate of Σ*Q* neurons predicts reaction time (Σ*Q*^+^: *n* = 0neurons,.*r*, = −0.97, *p* = 5.32 ×10^−6^; Σ*Q*^−^: *n* = 87 neurons, *r* = 0.84, *p* = 0.0026). Points show mean ± SEM (*n* = 9 mice). (**j**) Probability of switching choices decreases with increasing Σ*Q* (*r* = −0.94, *p* = 4.82 ×10^−5^). Points show mean ± SEM (*n* = 9 mice). (**k**) Pre-cue firing rate of Σ*Q* neurons predicts switching probability (Σ*Q*^+^: *n* = 60 neurons, *r* = −0.93, *p* = 1.06 ×10^−4^; Σ*Q*^−^: *n* = 87 neurons, *r* = 0.94, *p* = 5.91 ×10^−5^). Points show mean ± SEM (*n* = 9).

A core prediction of RL models is that rewarded trials increment Σ*Q* whereas unrewarded trials decrement it, with the magnitude of the updates shaped by the prior history of feedback events. We therefore tested whether claustrum neurons tracked these trial-by-trial Σ*Q* updates across the four possible outcome transitions between consecutive trials. Changes in firing rates closely tracked the direction and relative magnitude of the model-predicted updates for neurons whose activity scaled significantly positively (*n* = 60 of 341 neurons) or negatively (*n* = 87 of 341 neurons) with Σ*Q* (Fig. 3g). Thus, the activity of claustrum neurons tracks trial-to-trial adjustments of total value.

Total value (Σ*Q*) correlations were invariant to the choice of value-space axes. Fitting neuronal activity with a model including Σ*Q* and Δ*Q* instead of *Q*_*l*_ and *Q*_*r*_ identified a similar fraction of Σ*Q* neurons (Fig. S3d; 46%, *n* = 157 of 341 neurons), with 93% overlap between model formulations (Fig. S3e). Because Σ*Q* evolves slowly across trials, apparent neural correlations could arise from intrinsic firing rate autocorrelations or behavioral structure. To control for spurious correlations, we generated both Amplitude-Adjusted Fourier Transform (AAFT) surrogate neural data (Shin et al., 2021) that preserved each neuron’s autocorrelation structure while disrupting its relationship to Σ*Q* (Fig. S3f,g), as well as behaviorally matched pseudosessions generated by simulating choice and outcome sequences from mouse-specific *Q*-learning parameters (Harris, 2020), preserving reward statistics while eliminating learned value structure (Fig. S3h,i). After applying both surrogate controls, more than 85% of neurons originally classified as Σ*Q* neurons remained significantly correlated with Σ*Q* (Fig. S3j; *n* = 134 of 157 neurons using Σ*Q* and Δ*Q* model; *n* = 129 of 147 neurons using *Q*_*l*_ and *Q*_*r*_ model), indicating that Σ*Q* signals were not accounted for by firing rate drift or behavioral autocorrelation.

Claustrum neuron activity is modulated by licking and other movements (Shima et al., 1996; Ollerenshaw et al., 2021; Chevée et al., 2022). Furthermore, recent work has shown that claustrum activity is linked to behavioral state (Atlan et al., 2018; Fodoulian et al., 2020; Atlan et al., 2024), raising the possibility that apparent Σ*Q* signals reflect broad behavioral state fluctuations. We therefore first extracted face and body motion energy as measures of movement from synchronized behavioral videos (Fig. S4a). As face motion energy was more strongly correlated with Σ*Q* (Fig. S4b; Face motion energy: *r* = 0.24 ± 0.01; Body motion energy: *r* = 0.10 ± 0.01), we used face motion energy as a representative measure of movement. We also extracted pupil size as a proxy for behavioral state (Hess and Polt, 1964; Hoeks and Levelt, 1993). Including face motion energy alone, or together with pupil size, as additional regressors in the model did not significantly shift the distribution of Σ*Q t*-statistics relative to the base model (Fig. S4c; two-sample Kolmogorov-Smirnov (KS) tests: base model vs. base model + movement, *p* = 0.61; base model vs. base model + movement & pupil size, *p* = 0.4). Correspondingly, the fraction of neurons significantly correlated with Σ*Q* was largely preserved, with more than 85% retained under both models (Fig. S4d).

Total value (Σ*Q*) provides an estimate of reward availability in the environment, which has been linked to the regulation of behavioral vigor (Niv et al., 2007; Hamid et al., 2016; Bari et al., 2019) and exploratory behavior (Constantino and Daw, 2015; Wittmann et al., 2020; Zid et al., 2024). We therefore asked whether trial-by-trial variations in claustrum neuron firing rates covaried with behavioral readouts of foraging behavior. Reaction time decreased monotonically with increasing Σ*Q* (Fig. 3h; *r* = −0.93, *p* = 1.2 ×10^−4^). Similarly, for neurons with ITI firing rates positively correlated with Σ*Q*, reaction time decreased monotonically with increasing neural activity (Fig. 3i, Σ*Q*^+^: black; *r* = −0.97, *p* = 5.32 ×10^−6^). For neurons with ITI firing rates negatively correlated with Σ*Q*, reaction time increased monotonically with increased neural activity (Fig. 3i, Σ*Q*^−^: gray; *r* = 0.84, *p* = 0.0026). Similar relationships were observed for choice updating. The probability of switching choices decreased monotonically with increasing Σ*Q* (Fig. 3j; *r* = −0.94, *p* = 4.82 ×10^−5^). Similarly, for claustrum neurons with ITI firing rates positively correlated with Σ*Q*, switching probability decreased with increasing neural activity (Fig. 3k, Σ*Q*^+^: black; *r* = −0.93, *p* = 1.06*×*10^−4^), while for claustrum neurons with ITI firing rates negatively correlated with Σ*Q*, switching probability increased with increasing activity (Fig. 3k, Σ*Q*^−^: gray; *r* = 0.94, *p* = 5.91 ×10^−5^). These results indicate that claustrum neuron activity covaries with trial-by-trial adjustments in movement vigor and choice policy. Together, these findings are consistent with the conclusion that sustained claustrum activity reflecting Σ*Q* signals is not explained by action selection, measured movement during the ITIs, or arousal, and that this activity covaries with trial-by-trial foraging behavior.

### Individual claustrum neurons maintain stable representations of total value between trials

A central feature of RL frameworks is that value estimates are updated following feedback events and remain stable in the absence of new information, and thus predict that animals must maintain value representations between trials. Consistent with this prediction, individual claustrum neurons exhibited sustained firing rate changes reflecting Σ*Q* for at least 10 s across the ITIs (Fig. 4a–e). To quantify the representational stability of individual Σ*Q* neurons, we performed cross-temporal decoding (CTD) (King and Dehaene, 2014), training a linear discriminant analysis (LDA) decoder to classify trials with high versus low Σ*Q* using neural activity during the 1-s lick-free window immediately preceding the upcoming trial and testing classification accuracy on 1-s time bins across the subsequent trial. Claustrum neurons showed sustained above-chance decoding extending many seconds across the ITIs (Fig. 4c), demonstrating stable Σ*Q* representations during periods when no new task information is available. At the population level, firing rates of Σ*Q* claustrum neurons showed graded separation across Σ*Q* quintiles that stabilized shortly after task execution and persisted throughout the ITI (Fig. 4d,e).

**Figure 4.**
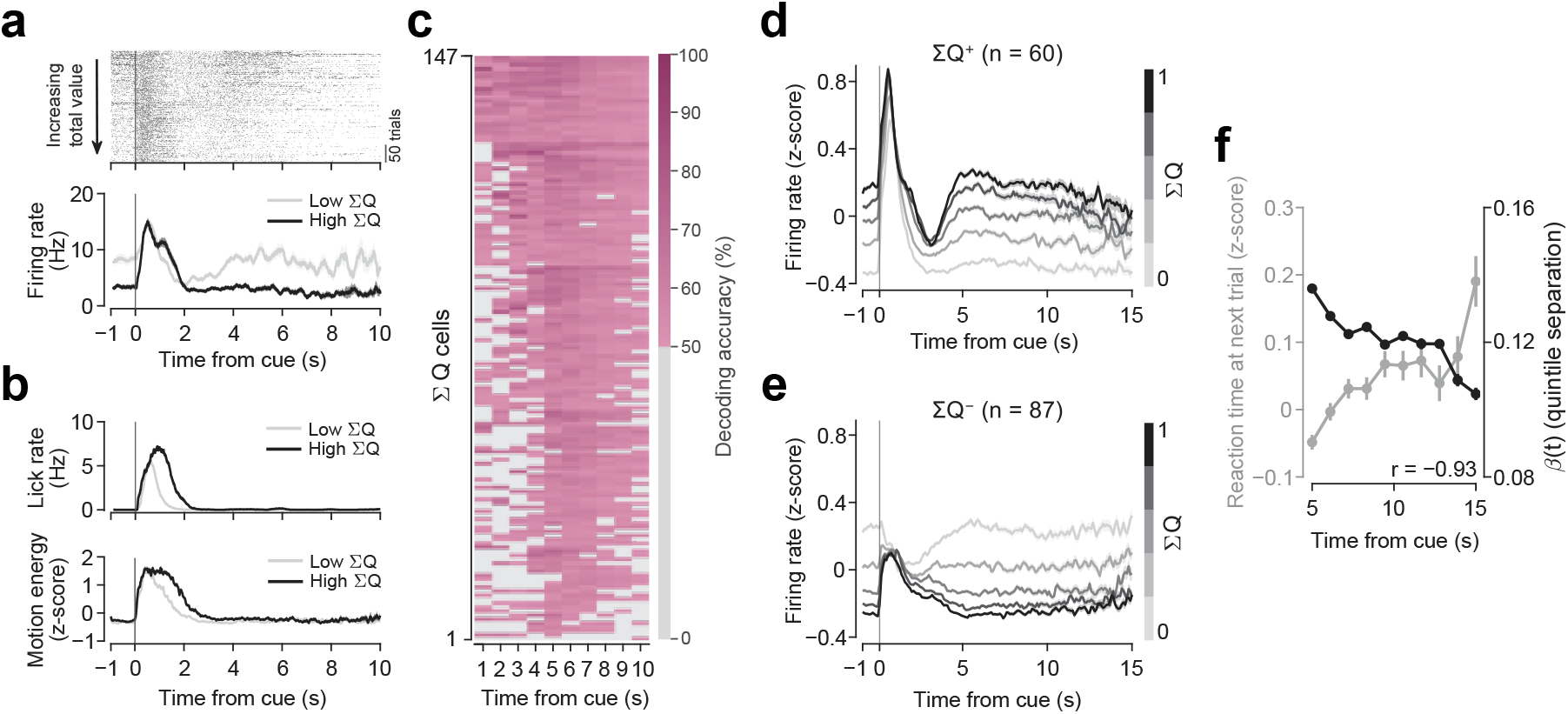
Stable total-value representations in individual claustrum neurons. (**a**) Example neuron showing sustained firing rate differences reflecting Σ*Q*. Top, cue-aligned raster plot with trials sorted by increasing Σ*Q*. Bottom, PSTHs for low (light) and high (dark) Σ*Q* trials (median split). (**b**) Lick rate (top) and face motion energy (bottom) aligned to cue onset for low (light) and high (dark) Σ*Q* trials for the neuron in (a). (**c**) CTD stability map for Σ*Q* neurons (*n* = 147*/*341 neurons). Rows denote neurons sorted by stability, columns denote time bins. Gray bins indicate chance-level decoding; colored bins indicate significant decoding. Higher color saturation indicates higher decoding accuracy. (**d**) Population PSTHs for neurons whose activity is significantly positively correlated with Σ*Q* (Σ*Q*^+^; *n* = 60 neurons). *Z*-scored firing rates are plotted across Σ*Q* quintiles (light to dark); shading indicates SEM. (**e**) Same as in (d) for neurons whose activity is significantly negatively correlated with Σ*Q* (Σ*Q*^−^; *n* = 87 neurons). (**f**) Reaction time on the upcoming trial (gray, left axis) and Σ*Q* stability (*β*(*t*); black, right axis) are plotted as a function of intertrial length. *β*(*t*) and reaction time are negatively correlated (*r* = −0.93).

Prior studies have shown that behavioral vigor and reaction time scale with the elapsed time between trials (Fecteau and Munoz, 2003; Guitart-Masip et al., 2011; Bari et al., 2019; Oda et al., 2021). We observed that longer ITIs were associated with slower reaction times on the subsequent trial (linear slope = 0.16, *p* = 0.002, *n* = 119 sessions). Total value (Σ*Q*) representations in the claustrum decayed across the ITI, with the separation between Σ*Q* quintiles progressively decreasing (linear slope, *β*(*t*) = −2.7 ×10^−4^, *p* = 1.2 ×10^−27^, *n* = 147 neurons). The magnitude of this decay was strongly and negatively correlated with reaction time (Fig. 4f; *r* = −0.93). Together, these results show that individual claustrum neurons maintain stable, graded representations of Σ*Q* across many seconds during the ITI, and that the gradual decay of their sustained activity predicts behavioral slowing.

### Action potential properties distinguish two electrophysiological classes of claustrumneurons

As anatomical studies and *in vitro* recordings have identified different neuronal cell types in the claustrum (Kim et al., 2016; Graf et al., 2020; Erwin et al., 2021; Lei et al., 2025; Shelton et al., 2025; Fodoulian et al., 2025; Graf et al., 2026), we asked whether the electrophysiological properties of recorded neurons distinguished different cell types. We found that the extracellular waveforms of recorded claustrum neurons exhibited a bimodal distribution of mean waveform widths (Fig. 5a; Narrow-spiking (NS) neurons, mean: 0.22 ± 0.06 ms, *n* = 143 of 341 neurons; Wide-spiking (WS) neurons, mean: 0.61 ± 0.07 ms, *n* = 198 of 341 neurons). Although in neocortical and hippocampal recordings wide-spiking (WS) waveforms are typically attributed to excitatory projection neurons and narrow-spiking (NS) waveforms to fast-spiking inhibitory interneurons (Mountcastle et al., 1969; Csicsvari et al., 1998), some cortical and subcortical projection neurons have NS waveforms (Vigneswaran et al., 2011; Ono et al., 2017; Kisner et al., 2018; Zemel et al., 2023). A recent study reported that some claustrocortical neurons were parvalbumin-positive (PV) neurons with NS action potentials (Shelton et al., 2025).

**Figure 5.**
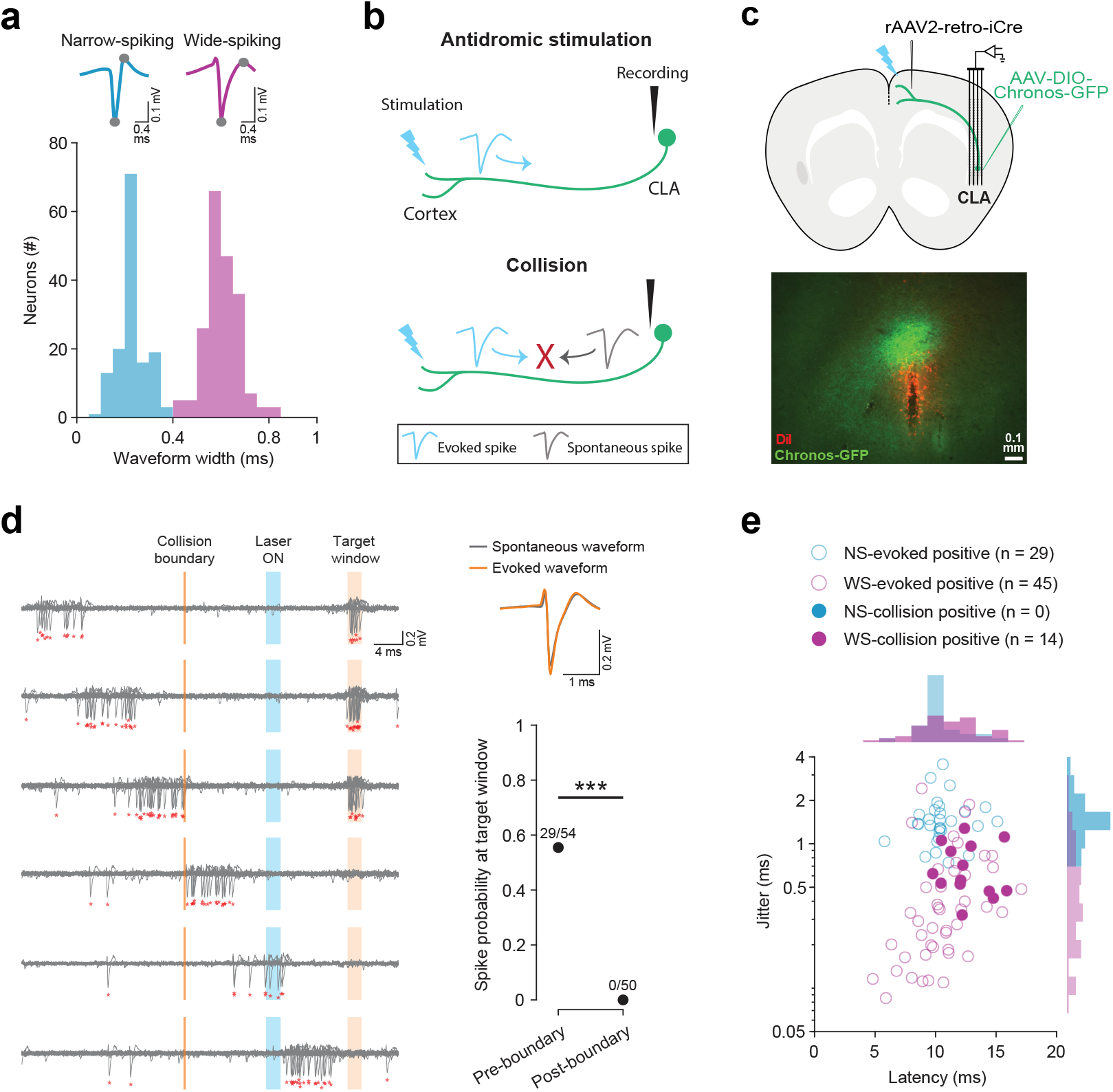
Wide-spiking (WS) neurons project to the medial frontal cortex. (**a**) Top: example narrow-spiking (NS) and wide-spiking (WS) waveforms recorded from claustrum neurons. Gray dots mark the valley and peak used to compute waveform width. Bottom, distribution of waveform widths across all claustrum neurons recorded during behavior (*n* = 341 neurons, *N* = 9 mice). (**b**) Collision test schematic. Spikes recorded in the claustrum evoked by cortical photostimulation (top). For collision-positive neurons, light-evoked spikes fail to propagate when a spontaneous spike is already traveling down the same axon, and are therefore not detected in claustrum recordings, verifying that the recorded neuron projects to the cortex. (**c**) Top, experimental configuration. rAAV2-retro-iCre was injected into ACC and RSC of C57BL/6J mice; AAV2-FLEX-Chronos-GFP was injected into the right claustrum. Neuropixel probes were inserted through the right claustrum. Bottom, coronal section centered on the claustrum showing Chronos-GFP expression (green) and Neuropixel probe tracks (red). Scale bar: 0.1 mm. (**d**) Example collision-positive neuron. Left, all traces with spontaneous action potentials in progressive 6 ms windows are plotted. Orange block indicates the time window when spikes evoked by cortical photostimulation are detected. Blue block indicates when the laser stimulation is on. Orange line indicates the calculated collision boundary. Right (top), average spontaneous (gray) and evoked (orange) spike waveforms (*r* = 0.99) for the example neuron. Right (bottom), evoked spike probability following spontaneous spikes before and after the collision boundary (Fisher’s exact test, *p* = 3.7 ×10^−11^). (**e**) Response latency versus the jitter of evoked action potentials for WS (Closed magenta circles: collision-positive, *n* = 14*/*59 neurons; Open magenta circles: collision-negative, *n* = 45*/*59 neurons) and NS neurons (Open blue circles: collision-negative, *n* = 0*/*29 neurons). WS neurons showed significantly lower jitter than NS neurons (Wilcoxon rank-sum, *p* = 1.5 ×10^−10^), while latency distributions did not differ (*p* = 0.19). Collision-positive neurons were significantly enriched among WS cells (Fisher’s exact test, *p* = 0.0037).

To test whether either of the two electrophysiological types project to frontal cortex, we performed antidromic collision testing (Fuller and Schlag, 1976; Li et al., 2015; Saiki et al., 2018), photostimulating claustrocortical axons expressing the excitatory opsin Chronos in medial frontal cortex while recording from claustrum neurons with a Neuropixels probe in an additional seven mice (Fig. 5b,c). We identified both WS and NS neurons (Fig. S5a), of which 29% were reliably activated with photostimulation (response reliability *>* 0.1; *n* = 88 of 300 neurons, 59 WS and 29 NS neurons; *N* = 7 mice). Collision-positive neurons exhibited a significant decrease in the probability of detecting light-evoked spikes within the target window following spontaneous spikes within the collision window, as seen in an example collision-positive neuron (Fig. 5d, Fig. S5b, top). A decrease in light-evoked spike probability in the target window was not observed for collision-negative neurons (Fig. S5b, bottom). Fourteen of the 59 WS neurons were collision-positive while none of the 29 NS neurons passed the collision test (Fig. 5e; Fisher’s exact test, *p* = 0.0037). In addition, we assessed the probability of detecting collisions and found that 21 of the 59 WS neurons lacked sufficient spontaneous action potentials to be statistically evaluated while all NS cells exhibited sufficient spontaneous action potentials for statistical assessment (Fig. S5c). Thus, 37% of testable WS neurons were collision-positive compared to 0% of NS neurons. In addition, although the response latencies did not differ between the two groups (Fig. 5e; WS: mean 11.1 ± 2.6 ms; NS: mean 10.5 ± 1.8 ms; Wilcoxon rank-sum test, *p* = 0.19), WS neurons exhibited significantly lower trial-to-trial jitter in their light-evoked responses as compared to NS neurons (Fig. 5e; WS: mean 0.6 ± 0.5 ms; NS: mean 1.5 ± 0.6 ms; Wilcoxon rank-sum test, *p* = 1.5 ×10^−10^), consistent with higher temporal fidelity of back-propagating action potentials along the axons of projecting neurons (Swadlow, 1998).

To further confirm our results, we immunostained claustrocortical neurons retrogradely labeled from frontal and midline cortical areas for PV expression, typically associated with a NS phenotype (Kim et al., 2016; White et al., 2018; Graf et al., 2020). Retrograde tracer injections into ACC, RSC and frontal motor areas labeled large populations of claustrocortical neurons. Across our experiments, 0.2% of the retrogradely labeled claustrocortical neurons were annotated as PV-positive (Fig. S5d–g; ACC and RSC: *n* = 8 of 3501 retrogradely labeled neurons; frontal motor areas: *n* = 1 of 227 retrogradely labeled neurons), representing 1% of PV-positive neurons (ACC and RSC: *n* = 8 of 731 PV neurons; frontal motor areas: *n* = 1 of 61 PV neurons). Overall, less than 1% of retrogradely labeled claustrocortical neurons expressed any inhibitory marker (ACC and RSC: *n* = 20 of 3501 neurons; frontal motor areas: *n* = 1 of 227 neurons). Together, these results indicate that claustrocortical neurons projecting to frontal and midline cortical areas are overwhelmingly excitatory neurons that do not express PV, consistent with our electrophysiological findings that claustrocortical neurons targeting frontal cortex are WS and not NS neurons.

### Narrow-spiking (NS) and wide-spiking (WS) neurons differ in their response properties during task execution

We next asked whether NS and WS neurons had distinct response profiles during task execution. The activity of almost all NS and WS neurons was significantly modulated relative to baseline following the go cue (Fig. 6a; NS: 94%, *n* = 134 of 143 neurons; WS: 86%, *n* = 171 of 198 neurons; Wilcoxon signed-rank test). However, the responses of NS neurons were significantly biased toward excitation (Fig. 6b, left; Excited: *n* = 97 of 143 NS neurons; Inhibited: *n* = 37 of 143 NS neurons; binomial test, *p* = 2.19 ×10^−7^), whereas the responses of WS neurons were significantly biased toward suppression (Fig. 6b, right; Excited: *n* = 51 of 198 WS neurons; Inhibited: *n* = 120 of 198 WS neurons; binomial test, *p* = 1.36 ×10^−7^), indicating that the responses of NS and WS neurons significantly differed during the animals’ responses following the go cue (Chi-squared test, *χ*^2^ = 54.49, *p* = 1.56 ×10^−13^). The distribution of directional preferences did not differ between classes (Chi-squared test, *χ*^2^ = 0.28, *p* = 0.60), with the responses of neither NS nor WS neurons having a significant bias for ipsilateral or contralateral choices (Fig. 6c,d; NS: Ipsilateral: *n* = 45 of 143 neurons, Contralateral: *n* = 42 of 143 neurons, binomial test, *p* = 0.83; WS: Ipsilateral: *n* = 43 of 198 neurons, Contralateral: *n* = 34 of 198 neurons, binomial test, *p* = 0.36). In contrast, the distribution of outcome preferences differed significantly between classes (Chi-squared test, *χ*^2^ = 6.14, *p* = 0.013). The responses of NS neurons showed a significant bias toward increased firing on rewarded relative to unrewarded trials, whereas WS neurons showed no such bias (Fig. 6e,f; NS: Rewarded: *n* = 62 of 143 neurons, Unrewarded: *n* = 39 of 143 neurons, binomial test, *p* = 0.028; WS: Rewarded: *n* = 42 of 198 neurons; Unrewarded: *n* = 54 of 198 neurons, binomial test, *p* = 0.26). Together, these results show that both NS and WS claustrum neurons are engaged by task events, yet differ in their response properties: NS neurons are predominantly excited and respond more strongly on rewarded trials, whereas WS neurons are predominantly suppressed and show no response bias to outcome.

**Figure 6.**
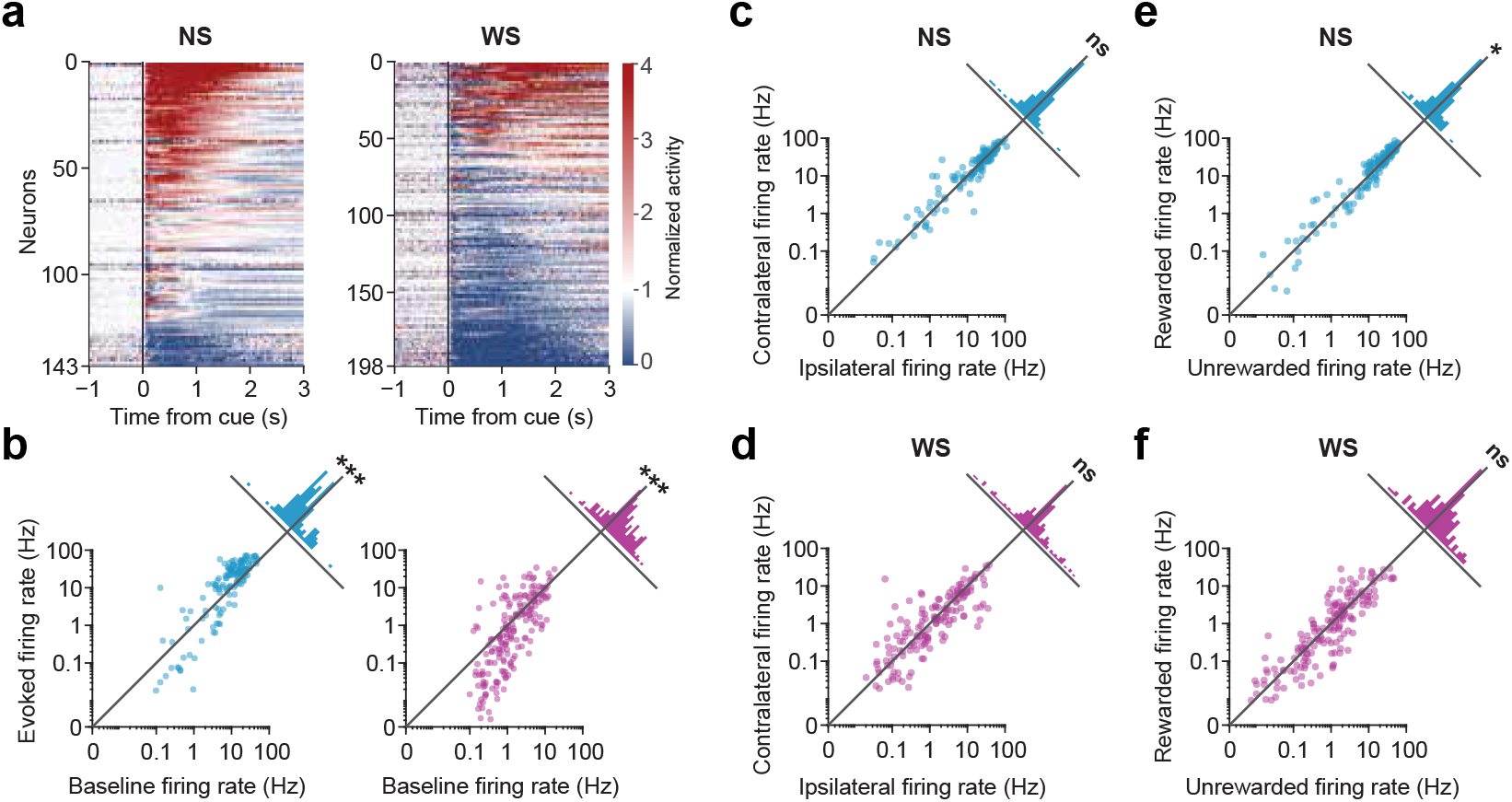
Responses of narrow-spiking (NS) and wide-spiking (WS) claustrum neurons during task execution. (**a**) Heatmaps of firing rates normalized to the 1-s pre-cue baseline for NS (*n* = 143) and WS (*n* = 198) neurons, aligned to cue onset. Neurons are sorted by their mean post-cue response. (**b**) Baseline versus task-evoked firing rates for NS (left) and WS (right) neurons. Each point represents one neuron. Diagonal lines indicate unity. Rotated marginal histograms depict the distribution of task-evoked changes in firing rate (ΔFR). NS neurons showed a significant bias toward excitation during task execution (NS Excited: *n* = 97*/*143 neurons; NS Inhibited: *n* = 37*/*143 neurons; binomial test, *p* = 2.19 ×10^−7^), whereas WS neurons were significantly biased toward inhibition during task execution (WS Excited: *n* = 51*/*198 neurons; WS Inhibited: *n* = 120*/*198 neurons; binomial test, *p* = 1.36 ×10^−7^). (**c**) Ipsilateral versus contralateral choice firing rates for NS neurons. Rotated marginal histograms depict ΔFR distributions. There was no significant bias for ipsilateral- or contralateral-choice-preferring NS neurons (Ipsilateral: *n* = 45*/*143 neurons; Contralateral: *n* = 42*/*143 neurons; binomial test, *p* = 0.83). (**d**) As in (c), but for WS neurons. There was no significant bias for ipsilateral- or contralateral-choice-preferring WS neurons (Ipsilateral: *n* = 43*/*198 neurons; Contralateral: *n* = 34*/*198 neurons; binomial test, *p* = 0.36). (**e**) Rewarded versus unrewarded firing rates for NS neurons. Rotated marginal histograms depict ΔFR distributions. NS neurons were enriched for neurons significantly modulated by rewarded trials (Rewarded: *n* = 62*/*143 neurons; Unrewarded: *n* = 39*/*143 neurons; binomial test, *p* = 0.028). (**f**) As in (e), but for WS neurons. There was no significant bias for reward-preferring or no reward-preferring WS neurons (Rewarded: *n* = 42*/*198 neurons; Unrewarded: *n* = 54*/*198 neurons; binomial test, *p* = 0.26).

### The firing rate of wide-spiking (WS) claustrum neurons during the intertrial intervals (ITIs) increases with decreasing total value

We next asked whether NS and WS neurons differed in their activity during the ITI, when claustrum activity correlates with Σ*Q*. Both NS and WS neurons exhibited sustained Σ*Q* representations across ITIs with comparable decay rates (Fig. S6a–f; Linear model: ITI length ×cell type, *t* = −0.01, *p* = 0.99). H., owever, the NS population was significantly enriched for neurons whose activity scaled with Σ*Q* relative to WS neurons (NS: 50%, *n* = 72 of 143 neurons; WS: 37%, *n* = 74 of 198 neurons; Fisher’s exact test, *p* = 0.02). In addition, NS neurons showed no population-level bias in the direction of Σ*Q* correlation: the number of NS neurons with firing rates scaling positively or negatively with Σ*Q* were not significantly different (Fig. 7a,b; binomial test, *p* = 0.40), and their value-space projections based on regression coefficients of *Q*_*l*_ and *Q*_*r*_ were indistinguishable from uniform (Rayleigh test, *p* = 0.38; mean resultant *R* = 0.08; *θ* = 19.5^*◦*^), consistent with no bias along the Σ*Q* axis (Fig. 7c; Σ*Q*^+^: *n* = 41 of 72 neurons, 57% versus Σ*Q*^−^: *n* = 31 of 72 neurons, 43%; binomial test, *p* = 0.29). In contrast, WS neurons were significantly more likely to increase their firing rates with decreasing Σ*Q* (Fig. 7d,e; binomial test, *p* = 0.0035), and their angular distribution in value space was significantly non-uniform (Rayleigh test, *p* = 8.3 ×10^−6^), with a larger mean resultant length aligned with the negative Σ*Q* axis (*R* = 0.24; *θ* = 222.5^*◦*^). Consistent with this, the majority of WS neurons with significant Σ*Q* coefficients exhibited negative Σ*Q* correlations (Fig. 7f; Σ*Q*^+^: *n* = 19 of 74 neurons, 26% versus Σ*Q*^−^: *n* = 55 of 74 neurons, 74%; binomial test, *p* = 3.38 ×10^−5^). Together, these results show that, although both NS and WS neurons represent Σ*Q*, WS neurons largely scale their activity inversely with Σ*Q*. As this population includes claustrocortical neurons projecting to medial frontal cortex, our findings indicate that graded increases in claustrocortical input to medial frontal cortex signal decreasing Σ*Q*, moments associated with adjustments in movement vigor and choice switching (Fig. 3h-k).

**Figure 7.**
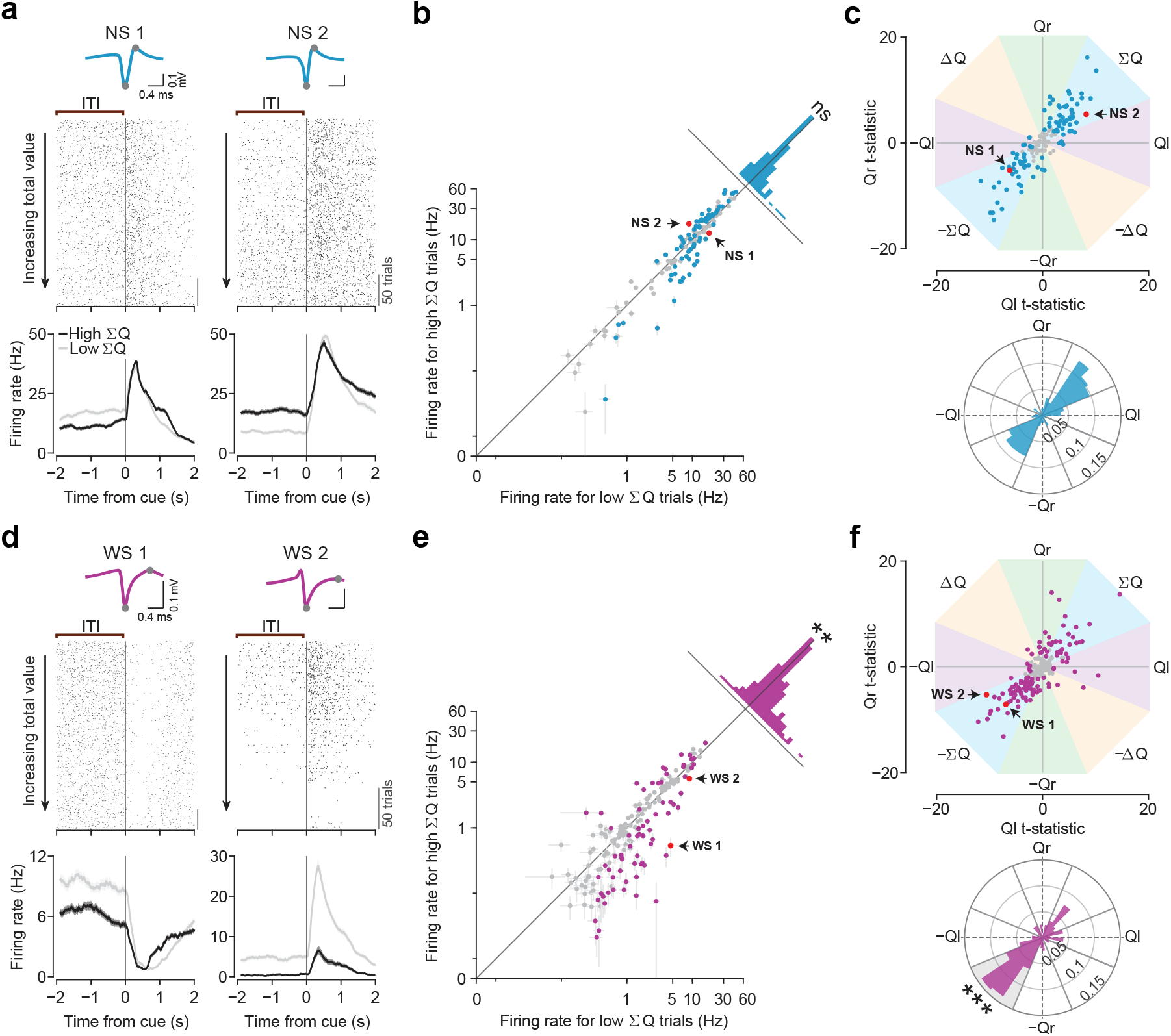
The firing rate of wide-spiking (WS) claustrum neurons during the intertrial intervals (ITIs) scales inversely with total value. (**a**) Example NS neurons. Top, spike waveforms. Middle, cue-aligned raster plots sorted by increasing Σ*Q*. Bottom, PSTHs for low (light) and high (dark) Σ*Q* trials (median split). (**b**) Scatter plot of firing rates for NS neurons during low versus high Σ*Q* trials. Cyan points indicate NS neurons significantly correlated with Σ*Q*; gray points indicate NS neurons with no significant correlation with Σ*Q*; red points indicate example neurons shown in (a). Rotated marginal histograms depict the distribution of changes in firing rate (ΔFR) between low and high Σ*Q* trials. (**c**) Value-space representation of NS neurons. Top, cyan points indicate neurons with significant value coefficients, gray points indicate the remaining NS neurons. Arrows mark example neurons from (a). Bottom, polar histogram of tuning angles, normalized as probabilities, and overlaid with sector boundaries. NS neurons showed no significant enrichment for positive versus negative Σ*Q* correlations (binomial test, *p* = 0.29). (**d**) Example WS neurons, plotted as in (a). (**e**) Scatter plot of firing rates for WS neurons during low versus high Σ*Q* trials, plotted as in (b). Arrows mark example neurons from (d). WS neurons were significantly biased toward higher activity in low Σ*Q* trials (binomial test, *p* = 0.0035; asterisks). (**f**) Value-space representation of WS neurons, plotted as in (c). Arrows mark example neurons from (d). WS neurons showed a significant enrichment for negative Σ*Q* correlations (binomial test, *p* = 3.38 ×10^−5^; asterisks).

## Discussion

During dynamic foraging, we found that most neurons recorded from anterior claustrum represented licking, choice, and outcome during task execution and that, between trials, almost half of these neurons represented trial-by-trial fluctuations in total value. Individual neurons sustained their activity over seconds across the intertrial intervals, consistent with RL predictions of stable value signals between trials, and the magnitude of these claustrum total-value signals predicted adjustments in reaction time and choice switching. We further identified two electrophysiologically distinct neuronal populations based on spike waveform. Using the spike-collision test (Fuller and Schlag, 1976; Li et al., 2015), we found that none of the NS neurons we tested were collision-positive, while many WS neurons were confirmed to project to medial frontal cortex. We found that NS neurons were excited during task execution, and their activity showed bidirectional correlations with total value. In contrast, WS neurons were generally suppressed during task execution, and their ITI firing rates increased as total value decreased. These results reveal distinct claustrum cell types associated with different value representations during dynamic decision making.

The lack of collision-positive NS neurons in our electrophysiological results is consistent with our anatomical data indicating that PV-positive NS inhibitory interneurons in the claustrum are not retrogradely labeled from ACC, RSC or frontal motor areas. In our recordings, NS neurons were sampled more frequently than predicted by anatomical estimates of PV-positive NS interneurons in the claustrum (Takahashi et al., 2023). One possibility is that narrow waveforms do not uniquely identify PV-positive NS interneurons, as narrow spikes have also been reported for other interneuron and projection neuron classes in other brain areas (Ma et al., 2006; Kvitsiani et al., 2013; Courcelles et al., 2024; Vigneswaran et al., 2011; Ono et al., 2017; Kisner et al., 2018; Zemel et al., 2023). Another possibility is that a much larger fraction of NS PV-positive interneurons is active during dynamic foraging than the fraction of claustrocortical neurons. This possibility is consistent with *in vitro* studies showing that corticoclaustral input strongly recruits PV-positive NS interneurons while weakly driving or suppressing claustrocortical projection neurons (Kim et al., 2016; White et al., 2018), as well as *in vivo* experiments showing that corticoclaustral neurons in lightly anesthetized mice activate NS neurons while suppressing ipsilateral WS claustrum neurons (Shaker et al., 2025). This circuit organization may also help explain differences in the response properties we identified between NS and WS neurons: NS neurons were predominantly excited whereas WS neurons were preferentially suppressed during task execution. How the circuit organization of NS and WS neurons contributes to the differences in their responses during the ITIs, and its relationship to other cell-type classification and circuit studies of the claustrum (White and Mathur, 2018; Graf et al., 2020; Erwin et al., 2021; Qadir et al., 2022; Shelton et al., 2025; Lei et al., 2025; Fodoulian et al., 2025; Graf et al., 2026) requires further study.

Claustrum neuron activity correlated with licking and lick direction during task execution, consistent with prior studies identifying movement responses in the claustrum (Shima et al., 1996; Ollerenshaw et al., 2021; Chevée et al., 2022). Here, we found that the activity of a subset of claustrum neurons also represented trial outcome during task execution. As previously observed using a sensory-selection task (Chevée et al., 2022), claustrum neurons exhibited anticipatory choice-related activity during the ITIs. By implementing a probabilistic reward task that dissociates reward rate from choice, we also revealed that recorded claustrum neurons were enriched for neurons representing total value between trials. Prior studies have shown that claustrum activity is modulated by task engagement, attention, and cognitive load (White et al., 2018; White and Mathur, 2018; Fodoulian et al., 2020; Atlan et al., 2024; Huang et al., 2025). In the dynamic foraging task used here, mice responded reliably to the go cue with very few missed trials during recording sessions, indicating that the total value signals we identified are unlikely to simply reflect lapses in behavior or disengaged states. In addition, in contrast with prior studies of claustrum neuron activity that reported elevated delay-period firing at the population level (Han et al., 2024; Bhattacharjee et al., 2024), our results indicate that individual claustrum neurons exhibit sustained activity correlated with total value for many seconds across ITIs.

Although claustrum activity during task execution exhibited lick-direction selectivity reminiscent of neurons in anterolateral motor cortex (ALM) (Guo et al., 2014; Li et al., 2015), ALM shows sparse value representations during the ITI in similar foraging tasks (Hattori et al., 2019; Bari et al., 2019), in contrast to our findings in the claustrum. The value signals represented in anterior claustrum also differ from those previously reported in other interconnected cortical regions studied using similar foraging paradigms, including medial prefrontal cortex (mPFC) (Bari et al., 2019), orbitofrontal cortex (Hattori et al., 2023), and RSC (Hattori et al., 2019). In these studies, neural activity typically represented mixtures of total value, relative value, chosen value, and action values, while we found that claustrum neurons were significantly enriched for total value representations. Given differences in task structure across experiments, it remains to be determined whether this representational profile is unique to the claustrum or would also be observed in other cortical regions under precisely matched task conditions.

Persistent neural activity plays a key role in flexible behaviors (Miller and Cohen, 2001; Wang, 2021). The cellular and circuit mechanisms that underlie the stability of claustrum total-value signals over extended timescales remain unknown. Intrinsic and circuit mechanisms (Major and Tank, 2004; Zylberberg and Strowbridge, 2017), potentially interacting with neuromodulatory inputs to the claustrum (Baizer, 2014; Zingg et al., 2018; Norimoto et al., 2020; Terem et al., 2020; Wong et al., 2021), may contribute to this sustained activity. Prior studies have demonstrated that recurrent thalamocortical circuits are necessary for sustaining preparatory activity and working memory in frontal areas (Guo et al., 2017b; Schmitt et al., 2017; Bolkan et al., 2017). For example, recordings revealed similarly selective persistent delay activity predicting movement direction in both ALM and its associated thalamic nuclei (Guo et al., 2017b), supporting theoretical models in which recurrent loops between areas with similar representations maintain persistent activity (Compte, 2000). The claustrum and frontal cortex are also highly reciprocally connected (Clascá et al., 1992; Smith and Alloway, 2010; Torgerson et al., 2015; Atlan et al., 2017; Wang et al., 2017; White et al., 2017; Zingg et al., 2018; Chia et al., 2020; Marriott et al., 2021; Wang et al., 2023), and recent work showed that corticoclaustral neurons in frontal cortex convey value signals to the claustrum. In a task in which mice assessed reward magnitude and reward probability, corticoclaustral neurons in dorsomedial frontal cortex encoded both expected value and reward risk (Majumdar et al., 2025), suggesting that reciprocal claustrocortical loops may contribute to the stability of value signals over extended timescales. Interestingly, circuit-mapping studies in frontal cortex and RSC indicate that claustrocortical and thalamocortical inputs preferentially activate different postsynaptic cell types, suggesting dissociable contributions to cortical representations (Jackson et al., 2018; Brennan et al., 2021; Anastasiades and Carter, 2021). How claustral and thalamic circuits interact to jointly shape cortical representations in frontal cortex remains an important open question.

Our finding that WS neurons monotonically increase their firing rate as total value declines provides a quantitative link between claustrum activity and flexible behavior. In this task, total value predicted response times, switching probability, and the progressive slowing of behavior across longer ITIs, collectively suggesting that total value functions as a global estimate of reward availability that regulates foraging behavior. This interpretation is consistent with frameworks in which animals maintain a running estimate of environmental richness that guides behavioral adjustments, promoting persistence when rewards are abundant and switching when they decline (Constantino and Daw, 2015; Wittmann et al., 2020; Zid et al., 2024), and regulating response vigor and latency (Niv et al., 2007; Guitart-Masip et al., 2011; Wang et al., 2013; Hamid et al., 2016; Bari et al., 2019). The prominence of total value representations in WS claustrum neurons suggests that the claustrum broadcasts this global estimate to cortical circuits involved in value-based decision making, thereby influencing the behavioral adjustments that foraging demands. Whether claustrum neurons exhibit similarly persistent representations of total value in tasks with different foraging demands remains an important direction for future work.

## Methods

### Animals

All procedures were approved by the Johns Hopkins Animal Care and Use Committee and followed the guidelines of the Society for Neuroscience and the National Institutes of Health. C57BL/6J mice (RRID: IMSR JAX:000664) were used for *in vivo* electrophysiology experiments. *Gad2-T2a-NLS-mCherry* mice (Peron et al., 2015) (RRID: IMSR JAX:023140) were used for anatomical experiments. Mice ranged in age from 1 to 6 months, and both males and females were used. Animals were kept on a 12 h light/dark cycle and provided with unlimited food and water until water restriction began.

### Surgery

Surgeries were performed on 6–12-week-old mice. Mice were initially anesthetized with isoflurane (3–5% in O_2_) for induction, followed by ketamine (50 mg/kg) and dexmedetomidine (25 *µ*g/kg), with anesthesia subsequently maintained with isoflurane (1.0–1.5% in O_2_). Mice were placed in a custom-built stereotaxic frame on a heated platform, and body temperature was monitored with a rectal probe. Ophthalmic ointment was applied to prevent corneal drying, and sterile saline (0.02 mL/g) and atropine (0.065 mg/kg) were administered subcutaneously. After disinfecting the scalp, an incision was made, and Bregma was used as the zero reference point. Buprenorphine (0.05 mg/kg, s.c.) was administered during or after surgery for analgesia. Mice recovered under a heating lamp before being returned to their home cages.

To selectively express Chronos in the claustrum, three small craniotomies were made over the left retrosplenial cortex (RSC) and anterior cingulate cortex (ACC). Coordinates are reported as AP/ML from Bregma and DV from the brain surface. Injections of rAAV2-retro-Syn-iCre (Janelia Research Campus Viral Tools Team) (Tervo et al., 2016) (60 nL per site) were made into posterior RSC (AP: −1.2 mm, ML: +0.3 mm, DV: −0.5 mm), anterior RSC (AP: −0.5 mm, ML: +0.3 mm, DV: −0.5 mm), and ACC (AP: +0.8 mm, ML: +0.5 mm, DV: −0.8 mm). In some mice, an additional injection targeted anterior ACC (AP: +1.5 mm, ML: +0.5 mm, DV: −0.8 to −1.2 mm). A second craniotomy was made over the claustrum, and AAV2-DIO-Chronos-GFP (University of North Carolina Vector Core) (Klapoetke et al., 2014) (120–240 nL) was injected at AP: +1.7 mm, ML: +2.6 mm, DV: −3.5 mm (from Bregma). Pipettes were left in place for 5–10 min before retraction. The skull was lightly roughened with a dental drill (Athena Champion AC5000, NTI Diamond F132-008 diamond burr), and treated with Liquid Dentin Activator for few seconds before attaching a custom-made titanium headplate using dental adhesive (C&B-Metabond, Parkell).

For behavioral recordings (*n* = 9 mice, 7 female and 2 male), a custom microdrive carrying 8 tetrodes and an optic fiber (Thorlabs, FT200UMT; 200 *µ*m core, 0.39 NA) was implanted during the same surgery. The implant was initially positioned above the claustrum at AP: +1.7 mm, ML: +2.6 mm from Bregma, DV: − 1.5 to −1.8 mm from the brain surface. Tetrode tips were positioned 0.5–1 mm deeper than the fiber tip. Tetrodes were coated with red retrobeads (Lumafluor) to facilitate post hoc track visualization (Chevée et al., 2022). A ground screw was implanted above the cerebellum and secured along with the microdrive to the headplate using dental cement (Jet Repair Acrylic Pound Package, Pearson Dental), and the remaining craniotomies were sealed with a thin layer of Kwik-Cast (World Precision Instruments). Mice were allowed at least 1 week to recover before water restriction. During water restriction, mice had *ad libitum* access to food and were monitored daily to maintain ∼85% of baseline body weight.

For collision-test experiments, a separate cohort of mice (*n* = 7, 6 female and 1 male) received the same viral injections described above targeting the right ACC, RSC, and claustrum, and recovered for at least three weeks. Prior to acute recordings, mice received dexamethasone (2–5 mg/kg, i.m.) to reduce brain swelling, a 1 ×1 mm craniotomy was opened over the claustrum (AP: +1 to +2 mm, ML: +2.2 to +3.2 mm), and a second 0.5 ×2 mm craniotomy was opened over the medial frontal cortex (AP: 0 to +2 mm, ML: 0 to +0.5 mm). Craniotomies were then covered with Kwik-Cast until the mouse recovered from anesthesia and rested for at least 3–4 hours in their home cage before the first acute recording session. Each mouse underwent one to three acute recording sessions for collision testing.

### Behavioral task

Mice were trained over three progressive stages to perform probabilistic value-guided foraging (Bari et al., 2019; Grossman et al., 2022; Su and Cohen, 2022).

#### Stage 1: habituation (days 1–2)

Before formal training, water-restricted mice were habituated to head fixation for 1–2 days while receiving free water from two lick spouts (21-gauge stainless steel tubes) positioned in front of a 38.1-mm diameter acrylic tube in which the mice rested. The spouts were mounted on a micromanipulator equipped with a manual linear encoder providing 10 *µ*m resolution (World Precision Instruments, M325). Licks were detected using a custom circuit (Janelia Research Campus, 2019-053) (Hayar et al., 2006). All task events were controlled and logged using custom Arduino code running on the same microcontroller platforms. Water rewards (2 *µ*L) were calibrated weekly. During this habituation period, mice learned to lick either spout to obtain water. Licks at any time produced water delivery according to reward probabilities drawn from the set {0, 1}, with the rewarded spout alternating every 20 trials.

#### Stage 2: structured blocks (days 3–10)

In the second training stage, the full trial structure was introduced. Each trial began with a 0.5 s auditory cue, which was a “go” cue on 90% of trials and a “no-go” cue on the remaining 10%. Auditory stimuli consisted of 7.5 kHz and 15 kHz square-wave tones, assigned as go and no-go cues, respectively. These tones were generated by microcontrollers (ATmega16U2 or ATmega328), amplified, and played through speakers (CUI Devices, GF0401M). Following a go cue, mice could lick either spout during a 1.7 s response window; licks triggered water delivery from the chosen spout. Reward probabilities initially alternated between {0, 1} for 1–2 days, after which probabilities were drawn from {0.1, 0.9} for the remainder of Stage 2. Licks following a no-go cue were neither rewarded nor punished. Block lengths were 20–35 trials. After the 3 s fixed trial duration, intertrial intervals (ITIs) were drawn from an exponential distribution with rate parameter 0.3 s^−1^, bounded between 0.1–20 s. This produced a flat hazard rate, ensuring that the likelihood of the next trial did not increase with ITI duration. The resulting mean ITI was 4.83 s. To minimize spontaneous licking, a 1 s no-lick window was introduced before tone onset. Licks within this window triggered a newly generated ITI followed by another 1 s no-lick window. This rule greatly reduced spontaneous licking. To dissociate outcome delivery from the first lick, a delay between choice and outcome was introduced: 100 ms initially, gradually increasing to 300 ms. During stage 2 only, small amounts of water were occasionally delivered manually to shape the animal’s behavior, encourage use of the response window, and promote switching. If a mouse developed a directional bias during a session, the spouts were shifted horizontally in the opposite direction prior to the next session.

#### Stage 3: final task structure with independent reward probabilities

In the final stage, each spout was assigned a reward probability drawn pseudorandomly from {0.1, 0.5, 0.9}. Probabilities were assigned to spouts independently, with block lengths drawn from a uniform distribution of 20–35 trials. To avoid synchronized block transitions, the first block of each session was staggered by drawing one spout’s block length from 6–21 trials. Probability values could not repeat on the same spout across consecutive blocks, and both spouts were never assigned 0.1 simultaneously to maintain task engagement. If one spout carried a reward probability greater than or equal to that of the other spout for three consecutive blocks, its probability was forced to 0.1 to encourage switching and prevent directional biases. To counteract severe perseveration, if a mouse chose the 0.1-probability spout for four consecutive trials, the next block for both spouts was extended by an additional 4 trials. This prevented mice from repeatedly choosing a low-value spout until its probability increased. Mice completed an average of 422 trials (range: 189–688) over 59 min (range: 29–89) per session.

### Electrophysiology and data pre-processing

In behaving mice, we recorded extracellular activity at 30 kHz using an Intan acquisition system (RHD2000 series multi-channel amplifier with an RHD2132 headstage). The system was connected to eight implanted tetrodes (32 total channels; nichrome wire, PX000004, Sandvik) fed through 39 ga polyimide guide tubes and mounted on a custom 3D-printed microdrive that allowed advancement of the tetrodes via a single screw. Tetrode impedances were reduced to 180–300 kΩ by gold plating. Each bundle of eight tetrodes was wrapped around a 200 *µ*m optic fiber used for optogenetic modulation of recorded units. After each recording session, the tetrode-fiber assembly was advanced 40– 75 *µ*m. Raw signals were median-subtracted across channels and high-pass filtered at 300 Hz using custom MATLAB software. Putative spikes were detected by thresholding the filtered signal *x* at 3*σ*_*n*_, where *σ*_*n*_ = median(|*x*| */*0.6745) (Quiroga et al., 2004). Detected events were sorted offline into single-unit clusters (SpikeSort3D, Neuralynx Inc.) based on waveform peak, trough, and principal components. Isolation quality was assessed using three criteria: few interspike interval violations (*<* 0.5% of total cluster spikes within a 2 ms interspike interval), a minimum waveform amplitude of at least 80 *µ*V, and an L-ratio below 0.05 (Schmitzer-Torbert et al., 2005). Units were additionally excluded if waveform drift was detected, quantified by computing waveform correlations in 5-min bins across the behavioral session; any unit whose correlation dropped below 0.9 between consecutive bins was considered unstable and removed from further analysis (Fig. S1f).

Electrophysiological recordings for collision-test experiments were performed with four-shank Neuropixels 2.0 probes (Steinmetz et al., 2021), using SpikeGLX (https://billkarsh.github.io/SpikeGLX/). Data were acquired at a sampling rate of 30 kHz and band-pass filtered between 0.3 and 6 kHz. Raw data were processed with the automated spike-sorting algorithm Kilosort 2.5 (Pachitariu et al., 2024), followed by manual curation in Phy (https://github.com/cortex-lab/phy). Single units were included in the data set as defined by: few interspike interval violations (*<* 0.5% of total cluster spikes within a 2 ms interspike interval), a minimum waveform amplitude of at least 50 *µ*V, a minimum waveform correlation of 0.9 across 5-min bins throughout the recording and clear separation in feature space during manual curation. Units that did not meet these criteria were removed from further analysis.

### Confirmation of claustrum targeting

For tetrode-based recordings, we used three methods to determine whether recorded neurons were located in the claustrum region: online optogenetic modulation of recorded neurons after each behavioral session, post hoc confirmation of tetrode tracks through the claustrum and post hoc analysis of the location of electrolytic lesions performed at the end of the last behavioral session (Chevée et al., 2022).

At the end of each behavioral recording session, we determined which recorded neurons were responsive to optogenetic stimulation to delineate the claustral region. The implanted optic fiber was connected to a 473 nm laser (Laserglow Technologies, LRS-0473-PFM-00100-03), and trains of 10 ms light pulses were delivered at 10 Hz. Light power at the fiber tip was approximately 10 mW, and pulse timing was gated by an optical shutter system (Uniblitz LS2S2T1 with VCM-D1 driver) under Arduino (Uno R3) control. To determine whether individual units were significantly driven by the light, we applied the SALT statistical framework (Kvitsiani et al., 2013). Neurons were classified as light-responsive if their first-spike latencies were modulated within a 10 ms post-pulse window (*p <* 0.05), and only units generating on average *>* 0.1 spikes within this window were considered as modulated by light stimulation. Using these criteria, 112 of 341 neurons were classified as light responsive (Fig. S1h,i). Non-light-responsive units were also assigned to the claustrum if either: 1) the unit was recorded simultaneously on the same tetrode as a light-responsive unit, or 2) the recorded unit was recorded between a more dorsal and a more ventral location at which light-responsive units were identified on the same tetrode.

After the final recording session, we made electrolytic lesions to mark the most ventral position of the electrodes. Mice were initially anesthetized with isoflurane (3–5% in O_2_) for induction, with anesthesia subsequently maintained with isoflurane (1.0–1.5% in O_2_), and a lesion was made by passing 20 *µ*A for 10 s through two channels of a single tetrode from which optogenetically activated units were recorded. Animals were perfused 16–24 hours later to allow the lesion site to become clearly visible. After being deeply anesthetized with isoflurane, mice were transcardially perfused with 0.01 M phosphate-buffered saline (PBS), followed by 4% paraformaldehyde (PFA) in 0.01 M PBS. Brains were removed and post-fixed in 4% PFA for 3 h at room temperature on a gentle shaker, and then rinsed three times in 0.01 M PBS (10 min per rinse). Coronal sections were cut on a vibratome (Leica VT-1000S) at a thickness of 60 *µ*m. Sections were mounted using Aqua Poly/Mount (Polysciences). Chronos-GFP expression and tetrode tracks labeled with red beads (Lumafluor) were imaged. Imaging was performed on a Zeiss LSM 510 or LSM 800 confocal microscope using 10×(0.3 NA), 20×(0.8 NA), or 40×(1.3 NA) objectives, or on a Keyence BZ-X710 epifluorescence microscope using a 4×objective.

Only datasets from mice in which: (i) Retrobead-labeled tetrode tracks traversed the claustrum identified via Chronos-GFP expression in claustrocortical neurons, (ii) the location of the electrolytic lesion was consistent with the inferred tetrode trajectory, and (iii) optogenetically modulated neurons were identified during recordings were included in the final analysis. In total, 341 claustrum neurons recorded from 9 mice met our inclusion criteria.

For Neuropixels recordings, claustrum localization was determined by identifying light-responsive recorded units combined with post hoc histological confirmation of probe tracks. Neurons were assigned to the claustrum if they were reliably activated following photostimulation (reliability *>* 0.1) or were recorded at depths bounded by more dorsal and more ventral locations containing light-responsive units on the same shank. After the final recording session, brain tissue was processed as described above, and Neuropixels tracks labeled with DiI (Thermo Fisher, D3911) or DiD (Thermo Fisher, D7757) were assessed. Datasets were included from mice in which: (i) Neuropixels probe tracks traversed the claustrum identified via Chronos-GFP expression in claustrocortical neurons and (ii) neurons activated following photostimulation were identified. In total, 300 claustrum neurons recorded from 7 mice met these inclusion criteria.

### Video recording

Eye, face and body movements were recorded using two monochrome CMOS cameras (Thorlabs, DCC1545M). Eye videos were acquired with a 1.0×telecentric lens (Edmund Optics, 58-430) to minimize geometric distortion, face and body videos were captured with a 3.5 mm lens (Navitar, NMV-4WA). The eye and facial regions were illuminated with an infrared LED lamp (Joyzan IR illuminator), and additional white LED illumination (Chanzon diode) was used to maintain the pupil diameter within the middle of its dynamic range. Video recordings were obtained from the left eye and from the right side of the face and body. Behavioral features were extracted with the open-source MATLAB GUI FaceMap (https://github.com/MouseLand/FaceMap) (Syeda et al., 2024). Pupil diameter was measured from the eye video. Mean absolute motion energy, computed as the average absolute difference in pixel intensity between consecutive frames, was calculated from user defined regions of interest that encompassed the face identity region including the nose, whiskers, and jaw, as well as a separate region covering the body.

### Collision test procedure for identification of claustrocortical neurons

To determine whether narrow-spiking (NS) and wide-spiking (WS) claustrum neurons project to the medial frontal cortex, we performed antidromic collision testing (Fuller and Schlag, 1976; Li et al., 2015; Saiki et al., 2018) in a cohort of naïve mice expressing Chronos in claustrocortical neurons. Chronos was expressed in claustrocortical neurons by injecting AAV2-DIO-Chronos-GFP in the claustrum and rAAV2-retro-Syn-iCre in ACC and RSC as described above. At least three weeks after viral injections, awake, head-fixed mice underwent acute electrophysiological recordings with a four-shank Neuropixels 2.0 probe. Before insertion, the tips of the probe were briefly dipped in DiI (Thermo Fisher, D3911) ethanol solution and allowed to dry. In animals with multiple penetrations across the anterior–posterior axis, we alternated between DiI and DiD (Thermo Fisher, D7757) to distinguish successive penetrations. The probe was lowered slowly through a 1 ×1 mm craniotomy over the anterior claustrum (coordinates from Bregma: AP: +1 to +2 mm, ML: +2.2 to +3.2 mm) to a final depth of −2.5 to −3.25 mm below the brain surface, moving at a speed of 2–3 *µ*m/s. Once at the target depth, the probe was retracted by 50–100 *µ*m to release tension within the tissue and left in place for 10–15 minutes before recording began. Once the probe was fully inserted, the brain was covered with a layer of 1.5% w/v agarose (Sigma, A9414). An Ag/AgCl grounding wire was connected to the recording probe and immersed in the agarose adjacent to the implanted ground screw. Optical stimulation was delivered via an optic fiber (Thorlabs, FT200UMT; 200 *µ*m core, 0.394 NA) positioned over medial frontal cortex (coordinates from Bregma: AP: 0 to +2 mm, ML: 0 to +0.5 mm) and coupled to a 473-nm diode-pumped solid-state laser (Laserglow). The fiber was coated with black nail polish to reduce scattered light and minimize optical artifacts induced in the recordings. Stimulation consisted of 800–1800 pulses, each 2 ms in duration, delivered at 2 Hz, with a light intensity of approximately 40 mW measured at the fiber tip. Chronos activation evoked short-latency, stimulus-locked spikes in a subset of claustrum neurons. Neurons were classified as claustrocortical projection neurons if they showed: (i) reliable antidromic activation with a minimum reliability of 0.1, and (ii) a collision test outcome in which spontaneous spikes eliminated the subsequent stimulus-locked spike (see below for statistical procedures).

### Retrograde tracing

Retrograde neuronal tracing using Alexa Fluor–conjugated cholera toxin subunit B (CTB) was performed in two additional cohorts of mice using the anesthesia and surgical preparation described above. In one cohort (*N* = 4 male *Gad2-T2a-NLS-mCherry* mice), small craniotomies were made over the ACC (AP: +0.8 mm, ML: +0.3 mm from Bregma, DV: −0.8 mm from brain surface), anterior RSC (AP: −0.4 mm, ML: +0.3 mm from Bregma, DV: −0.5 mm from brain surface), and posterior RSC (AP: −1.2 mm, ML: +0.3 mm from Bregma, DV: −0.5 mm from brain surface). In the second cohort (*N* = 4 *Gad2-T2a-NLS-mCherry* mice, 2 female and 2 male), craniotomies were made over frontal motor cortex (AP: +2.5 mm, ML: +1.0 mm from Bregma, DV: −0.5 mm from brain surface). A total of approximately 180 nL of CTB was injected per mouse using a glass pipette (12.5–35 *µ*m tip), which remained in place for 5–10 min before withdrawal. The scalp was sutured, and mice recovered under a warm heat lamp.

Mice were sacrificed 11 days after injection in the first cohort and 7 days after injection in the second cohort. Following deep anesthesia with isoflurane, mice were transcardially perfused with 0.01 M PBS, followed by 4% PFA in 0.01 M PBS. Brains were removed and post-fixed in 4% PFA for 3 h at room temperature on a gentle shaker, and then rinsed three times in 0.01 M PBS (10 min per rinse). Coronal sections were cut on a vibratome (Leica VT-1000S) at a thickness of 60 *µ*m. Every third section was collected for immunohistochemistry. Sections were first incubated in a blocking solution containing 3% normal donkey serum and 0.3% Triton X-100 in 0.01 M PBS for 1 h at room temperature. Primary antibodies were rabbit anti-parvalbumin (PV; Swant, PV27, 1:1000) and mouse anti-GAD67 (Millipore, MAB5406, 1:1000). Sections were incubated with primary antibodies, diluted in the blocking solution, overnight at 4^*◦*^C. After three washes in PBS (10 min each), sections were incubated with secondary antibodies for 1–2 h at room temperature on a shaker. Secondary antibodies included donkey anti-rabbit DyLight 405 (Jackson ImmunoResearch, 711-475-152, 1:1000) for PV and donkey anti-mouse Alexa Fluor 647 (Thermo Fisher, A31571, 1:1000) for GAD67. Following secondary incubation, sections were rinsed three additional times in PBS (10 min each) and mounted with Aqua Poly/Mount.

For cell counting, images were collected using a Zeiss LSM 800 confocal microscope with a 20×objective (0.8 N.A) using *z*-stacks acquired at 2 *µ*m intervals. Claustrum images were obtained sequentially from anterior to posterior regions. High-magnification images were obtained with a 63×objective lens (1.4 N.A). Cell counting was performed using the Cell Counter plugin in FIJI (ImageJ, NIH) (Schneider et al., 2012). The boundaries of the claustrum were defined based on parvalbumin (PV) immunostaining (Druga et al., 1993). A binary matrix was constructed to represent the co-expression of four markers: ‘CTB’, ‘PV’, ‘GAD65-mCherry’, and ‘GAD67’ across populations of labeled cells. All unique, non-empty intersections among marker sets were quantified, together with the total number of cells expressing each marker irrespective of co-expression. Absolute counts were normalized to the total number of cells and expressed as population ratios. In total, 7,486 cells were counted from the ACC/RSC cohort and 571 cells from the frontal motor cortex cohort. Population ratios were visualized using a MATLAB UpSet plot (https://de.mathworks.com/matlabcentral/fileexchange/123695-upset-plot).

### Quantification and statistical analysis

All analyses were performed using MATLAB, R, and Stan. Data are reported as mean ±SEM unless stated otherwise. All statistical tests were two-sided and evaluated at a significance level of *α* = 0.05, unless stated otherwise. No-go trials were excluded from all analyses and treated as part of the intertrial interval except for the analysis specifically examining responses during no-go trials.

### Descriptive models of behavior

To quantify how recent reinforcement influenced choice behavior, we fit logistic regression models to predict choice as a function of past rewarded and unrewarded outcomes. For each mouse, we fit the model:

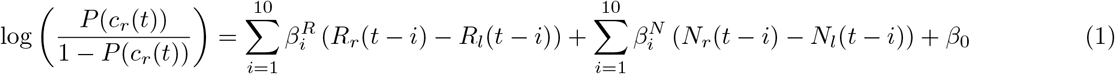

where *P* (*c*_*r*_(*t*)) denotes the probability of choosing the right option on trial *t, c*_*r*_(*t*) = 1 indicates a right choice and *c*_*r*_(*t*) = 0 indicates a left choice. *R*_*r*_ and *R*_*l*_ are binary variables indicating whether reward was delivered following a right or left choice on trial *t*, respectively (1 = rewarded, 0 = unrewarded). *N*_*r*_ and *N*_*l*_ are binary variables indicating whether a right or left choice was not rewarded on trial *t*, respectively (1 = unrewarded, 0 = rewarded). This model therefore quantifies how past rewarded and unrewarded outcomes on each side influence current choice.

To quantify the dependence of choice on past choices and rewards, we fit a second logistic regression model for each mouse:

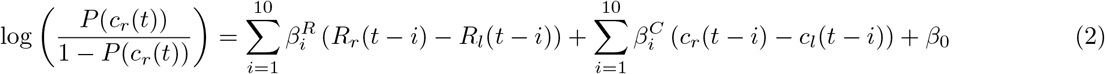

where *c*_*r*_(*t*) −*c*_*l*_(*t*) ∈ {−1, 1} denotes the animal’s choice on trial *t* (right = −1, left = −1), and *R*_*r*_(*t*) −*R*_*l*_(*t*) ∈ {−1, 0, 1} denotes the outcome on trial *t* (reward on the right = 1, reward on the left = −1, no reward = 0). This model therefore quantifies how past choices and rewards over the preceding 10 trials influence current choice.

### Integrated history estimates

To obtain trial-by-trial measures of choice and reward history, we derived scalar history estimates using exponential kernels fit to the regression coefficients from Eq. (2), independently for each behavioral session. We fit a single exponential function to the coefficients associated with the preceding ten trials:

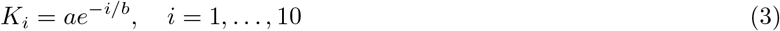

where *b* determines the decay timescale of the kernel and *a* is a scaling constant. The kernel was normalized so that 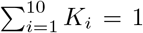, such that only the relative weighting across lags determines the history estimate. Separate kernels were derived for choice and reward history, denoted 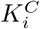 and 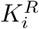.

Choice history was computed by convolving the choice sequence with the choice-history kernel and shifting it by one trial so that the value on trial *t* reflected only choices from trials *t* − 1 through *t* − 10:

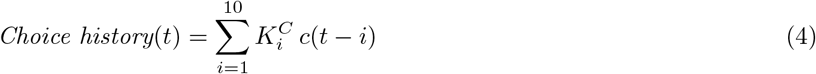

where 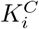 denotes the choice-history kernel at lag *i, c*(*t* − *i*) = *c*_*r*_(*t* − 1) −*c*_*l*_(*t* − 1) denotes choice on trial *t* − *i* (right = 1, left = −1). The resulting *Choice history* variable takes continuous values between −1 and 1.

Reward history was computed analogously using the reward-history kernel:

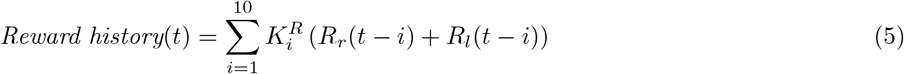

where 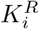 denotes the reward-history kernel at lag *i, R*_*r*_(*t* − *i*) and *R*_*l*_(*t* − *i*) are binary variables indicating whether reward was delivered following a right or left choice on trial *t* − *i*, respectively (1 = rewarded, 0 = unrewarded). Thus, the term *R*_*r*_(*t* − *i*) + *R*_*l*_(*t* − *i*) indicates whether reward was obtained on trial *t* − *i*, irrespective of choice side. The resulting *Reward history* variable takes continuous values between 0 and 1.

This approach yielded integrated choice and reward history estimates that reflect exponentially weighted recent behavior, matched to the empirically derived history timescales of each session.

### Generative models of behavior

We applied a family of reinforcement-learning (RL) models of behavior (Barraclough et al., 2004; Bari et al., 2019; Grossman et al., 2022; Su and Cohen, 2022). Q-learning estimates action values (*Q*_*l*_(*t*) and *Q*_*r*_(*t*)) on each trial, compares them, and generates choices. Choices are described by a random variable, *c*(*t*), corresponding to left or right choice, *c*(*t*) ∈ {*l, r*}. The value of a choice is updated as a function of the reward prediction error (*δ*), and the rate at which this learning occurs is controlled by the learning rate parameter (*α*). To capture asymmetric learning from positive and negative outcomes, we used separate learning rates for rewarded and unrewarded trials, denoted *α*_(+)_ and *α*_(−)_. In addition, the model included a forgetting parameter, (*ζ*), which allowed the value of the unchosen option to decay toward zero, representing increasing uncertainty about actions that have not been sampled. Thus, if the left option was selected on trial *t*, value updates followed:

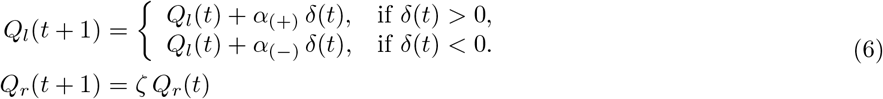

Choices were generated using a softmax decision function (Daw et al., 2006), which converts the difference in action values into a trial-by-trial probability of selecting each option:

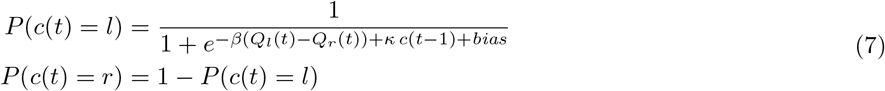

where *β* is the inverse-temperature, controlling the tradeoff between exploration and exploitation, *κ* captures choice autocorrelation, modeling the tendency to repeat the previous choice, and *bias* captures any consistent side preference independent of learned values.

This full model consistently outperformed reduced models lacking *κ* or *bias* in all animals (Fig. S3a). Improvement provided by *κ* was substantial, indicating perseverative behavior, whereas *bias* provided a smaller but reliable improvement, reflecting mild side biases. Because this model provided the best overall fit across animals, it was used for all subsequent analyses.

### Model fitting

We fit RL models (Q-learning) using Stan (Carpenter et al., 2017) through its R interface, RStan (https://mc-stan.org/rstan/), with all data preprocessing performed in R and MATLAB. We used a hierarchical Bayesian formulation in which all session-level parameters were modeled as draws from mouse-level (group-level) distributions. This partial pooling approach stabilized parameter estimation by reducing overfitting to noise in individual sessions while still capturing meaningful session-to-session variability in learning and choice behavior. Each session-specific parameter was modeled on an unconstrained latent scale as

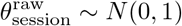

and then transformed into its appropriate constrained range using a probit transformation. On this latent scale, mouse-level group means *µ* and standard deviations *σ* were assigned weakly informative priors,

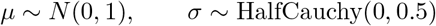

After transformation, the behavioral parameters were constrained to the following ranges: *α*_(+)_, *α*_(−)_ and *ζ* ∈ [0, 1], *β* ∈ [0, 10], *κ* ∈ [0, 5] and *bias* ∈ *R*. Sampling was performed using Stan’s Hamiltonian Monte Carlo No-U-Turn Sampler (NUTS). Five independent chains were run for 11,000 iterations each, including 5,000 warm-up iterations, yielding 30,000 post-warm-up samples. The target acceptance probability was set to 0.96 to ensure stable sampling with smaller step sizes. Default Stan tuning parameters were used unless otherwise specified.

### Extraction of model-derived variables and behavioral simulations

To extract latent model variables for analysis (i.e., trial-by-trial value estimates), we sampled 1,000 draws from the posterior distribution of the session-level parameters and, for each draw, ran the model forward through the actual sequence of choices and outcomes for that session. Model-derived variables (such as Q-values, relative value (Δ*Q*) and total value (Σ*Q*)) were then averaged across these posterior runs to obtain stable estimates of each latent quantity. To assess parameter estimates at the level of individual mice, we approximated each session’s maximum a posteriori (MAP) parameter value by estimating the mean of its posterior distribution (Fig. S3a). For behavioral simulations, we drew 1,000 samples from the posterior distribution of the mouse-level (group-level) parameters. For each draw, we simulated behavior in multiple synthetic sessions by generating choices and outcomes using the model’s choice rule and learning updates. This procedure allowed us to evaluate the model’s generative capacity and compare simulated behavior to the empirical data.

### Linear regression models of neural activity

To quantify how claustrum neuron activity correlated with behavioral variables, we fit linear regression models to trial-aligned, z-scored spike trains using MATLAB function “fitlm”. Time-resolved regressions were computed in 100-ms bins from the pre-cue period through the trial to identify neurons significantly modulated by lick rate, choice, or outcome. In addition, we performed regression analyses of predefined task epochs to quantify the fraction of neurons significantly modulated at behaviorally relevant times. Task execution was divided into four behavioral epochs: Epoch 1: 0–100 ms following the cue; Epoch 2: 0–0.3 s following the choice, the period between choice and outcome delivery; Epoch 3: 0.3–2 s following the choice, the period following outcome in which *>* 99% of licks occur during task execution; and Epoch 4: 2–3 s following the choice, when *<* 1% of licks occur during task execution. For each epoch, linear regression models tested whether neuronal firing rates were significantly explained by one or more task variables. To characterize value signals, ITI firing rates were regressed onto left action value (*Q*_*l*_), right action value (*Q*_*r*_) and choice. Each neuron’s regression coefficients were projected into a two-dimensional value space and classified into canonical subspaces (*Q*_*l*_, *Q*_*r*_, Σ*Q*, and Δ*Q*) based on polar angle (Seo et al., 2009; Wang et al., 2013). An equivalent formulation using Σ*Q* and Δ*Q* regressors was used to verify that classifications were not biased by axis choice. To control for movement and arousal, we extended the model to include video-derived motion energy and pupil diameter. Face and body motion energy were highly correlated (*r* = 0.64 ± 0.02), so face motion energy was used because it showed a stronger relationship with Σ*Q* (*r* = 0.24 ± 0.01). Analyses were restricted to behavioral sessions with available videos (Face: 89/119 sessions; Pupil: 71/119 sessions). Face motion energy was included as an additional regressor in a “+movement” model that could be fit for neurons recorded in sessions with face video (271/341 neurons), and pupil diameter was added in a “+movement & pupil” model that could be fit for neurons recorded in sessions with both videos (212/341 neurons). All regressors were z-scored, and significance was assessed at *α* = 0.05.

### Decoding of choice and outcome

To determine whether claustrum neurons encoded choice and outcome independently of licking and correlations between task variables, we compared decoding from raw firing rates to decoding from residual firing rates after removing variance explained by lick rate and the non-decoded task variable. For each neuron and task epoch, firing rates were residualized using a linear model that regressed out lick rate and outcome for choice decoding, and lick rate and choice for outcome decoding, such that residual activity reflected variance uniquely attributable to the variable of interest. We then performed single-neuron decoding using linear support vector machines (SVMs), using firing rates to predict either choice (ipsilateral vs. contralateral) or outcome (reward vs. no reward). Decoding performance was quantified as the area under the receiver operating characteristic curve (AUC) with 5-fold cross-validation. Statistical significance was assessed with a shuffle-null procedure in which trial labels were randomly permuted (50 shuffles per neuron and condition). A neuron was classified as informative if its true AUC exceeded the 95th percentile of the shuffle-null distribution.

### Nested model comparison using lagged regressors

To quantify the extent to which neural activity during the ITI reflects information from multiple past trials, we fit a series of nested linear models to the firing rate of each neuron. The base model (M1) included regressors for choice, outcome, and their interaction from the immediately preceding trial (*t*–1). We then constructed progressively more complex models by sequentially adding regressors from earlier trials (M2: *t*–1 and *t*–2; M3: *t*–1 to *t*–3; and so on, up to *t*–5). For each neuron, we compared successive models using likelihood ratio tests (LRTs) to determine whether the inclusion of additional lagged terms significantly improved model fit. The test statistic was computed as twice the difference in log-likelihood between the nested models and evaluated against a *χ*^2^ distribution, with degrees of freedom equal to the number of additional parameters. A model incorporating longer lags was considered to provide a significantly better fit if the LRT yielded *p <* 0.05. We quantified the fraction of neurons whose activity was better explained by models incorporating progressively longer lags, thereby estimating the temporal span over which past choice and outcome influenced neural firing.

### Comparison of Last-trial and History models

To determine whether claustrum neuron activity is better explained by integrated history estimates, we compared two regression models fit to ITI firing rates on a per-neuron basis. The Last-trial model included predictors for the choice and outcome of the immediately preceding trial (*t*–1) and took the form:

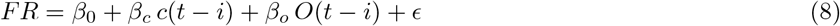

The History model replaced these predictors with scalar variables capturing exponentially weighted choice and reward history derived from behavioral fits (see “Integrated history estimates”), such that:

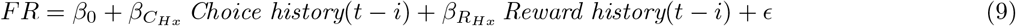

Both models were fit using linear regression to firing rates in the 1-s lick-free window preceding cue onset, and model performance was compared using Akaike Information Criterion (AIC; lower values indicate better fit). Each neuron was assigned to the model with lower AIC. After identifying neurons with activity better explained by the History model, we quantified the contribution of individual predictors using partial regression by comparing the full History model to reduced models in which either the choice-history or reward-history predictor was omitted, and computed the corresponding change in explained variance (Δ*R*^2^) to estimate the unique contribution of each term.

### Autocorrelation controls

To evaluate whether correlations between firing rates of claustrum neuron activity in the ITI and model-derived Σ*Q* reflected reliable task-related modulation rather than shared temporal autocorrelation, we performed two complementary surrogate-based control analyses. First, we generated amplitude-adjusted Fourier transform (AAFT) surrogates of the trial-by-trial pre-cue firing rates (Shin et al., 2021), which preserve the empirical autocorrelation and marginal distribution while randomizing the temporal phase. For each neuron, 500 surrogates were regressed onto latent variables, and the observed *t*-statistic was deemed significant if it exceeded the 95th percentile of the surrogate distribution. Second, to control for temporal structure intrinsic to the task and model-derived variables, we simulated behavior using the fitted reinforcement-learning model (Grossman et al., 2022; Harris, 2020). For each session, posterior samples of model parameters were used to generate 500 synthetic behavioral sessions of matched length. Latent variables were recomputed from these simulations by averaging across 100 posterior samples, and the real firing rates were regressed onto the simulated variables to obtain a null distribution of *t*-statistics reflecting correlations expected from task structure alone. A neuron was considered to significantly correlate with Σ*Q* only if its observed *t*-statistic exceeded both surrogate-based null distributions, providing a conservative control for autocorrelation in both neural and behavioral time series.

### Temporal stability of total value signals

To quantify the temporal stability of Σ*Q* signals at the single-neuron level, we performed cross-temporal decoding (CTD) (King and Dehaene, 2014) using trial-by-trial Σ*Q* estimates from the reinforcement-learning model. Spike trains were binned into non-overlapping 1-s windows aligned to cue onset. For each neuron, Σ*Q* values were binarized using a median split (high vs low Σ*Q*) to generate training labels. A linear discriminant analysis (LDA) classifier was trained on firing rates from the pre-cue 1-s bin using 10-fold cross-validation. The trained decoder was then tested on firing rates from all subsequent time bins (1–10 s), producing a cross-temporal decoding profile indexed by classification accuracy. To assess population-level stability of Σ*Q*, firing rates were aligned to cue onset and sign-aligned by each neuron’s Σ*Q* tuning. Trials were grouped into five Σ*Q* quintiles, and firing rates were averaged within each quintile and across neurons. At each time point, the strength of Σ*Q* representation was quantified as the slope *β*(*t*) of a linear fit across these quintile-averaged firing rates, providing a measure of how strongly population activity separated Σ*Q* representations over time. To test whether Σ*Q* representations evolved differently across NS or WS cell types, *β*(*t*) was modeled as a function of time, cell types, and their interaction. The main effect of time captured shared decay in Σ*Q* representation across the ITI, whereas the time ×class interaction quantified differences in decay rate between NS and WS populations.

### Spike collision testing

To determine whether a neuron was collision positive, we first required a minimum reliability for spikes evoked by photostimulation of ≥ 0.1 across laser pulses. For each neuron, we computed the distribution of light-evoked spike latencies and defined the back-propagation time as the median of this distribution. The antidromic target window was defined as the *Q*_10_–*Q*_90_ percentile range of the latency distribution, providing an estimate of the time window in which evoked spikes were reliably detected. Using the leading edge of this window (*Q*_10_) we defined a collision window during which a spontaneous orthodromic spike would be expected to collide with an antidromic spike traveling in the opposite direction. The collision window extended from

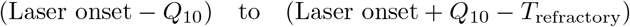

where *T*_refractory_ was set to 2 ms to enforce a physiologically plausible absolute refractory period. We compared the probability of detecting an antidromic spike within the target window following: (i) spontaneous spikes occurring inside the collision window, and (ii) spontaneous spikes occurring in an equally sized pre-collision control window immediately preceding the collision boundary. Fisher’s exact test with a more conservative alpha (*α <* 0.01) was used to determine whether spontaneous spikes significantly reduced the probability of antidromic activation. If *p <* 0.05 is used as a statistical cut-off, 3 additional WS neurons pass the collision test and no NS neurons pass. To assess the statistical feasibility of detecting collisions given observed spike counts, we performed a theoretical detectability analysis assuming idealized conditions in which spontaneous spikes occurring in the collision window always abolished light-evoked spikes, and each photostimulation evoked a spike when spontaneous spikes occurred before the collision window. For a range of pre- and post-boundary spike counts, Fisher’s exact test (*α* = 0.01) was used to identify neurons that were underpowered for collision detection based on spike counts alone.

### Waveform-based response properties

Waveform width for each neuron was computed as the number of recording samples between the waveform valley and the first subsequent sample at which the derivative changed sign, corresponding to the valley-to-peak duration. Neurons were classified as NS or WS using a threshold of 0.4 ms. For task-evoked analyses, firing rates were normalized relative to each neuron’s 1 s pre-cue baseline, and trial-averaged firing rate traces were used to generate population heatmaps. Ipsilateral versus contralateral choice modulation was assessed from −100 to 100 ms around the first lick using Wilcoxon signed-rank tests, and rewarded versus unrewarded outcome modulation was assessed from 0 to 500 ms after outcome delivery using the same test. Population-level biases were assessed using binomial tests on the fraction of significantly modulated neurons, and differences between NS and WS populations were evaluated using chi square tests.

## Acknowledgments

We thank Daeyeol Lee for discussions related to the project, Eun Ju Shin for sharing code related to the AAFT analysis, and Maxime Chevée, Marshall Hussain-Shuler, and members of the Brown laboratory for comments on the manuscript. This work was supported by a Kavli Neuroscience Discovery Institute Distinguished Graduate Student Fellowship (ABT), a Natural Sciences and Engineering Research Council of Canada PGD-D (SYA), the National Institute of Mental Health (R56 MH138416, SPB), and the National Institute of Neurological Disorders and Stroke (R01 NS085121, SPB; RF1 NS131984, JYC).

## Author contributions

ABT, JYC and SPB conceptualized the study. ABT, SYA, JYC and SPB designed the experiments. ABT, SYA and SJK performed the experiments. ABT, SYA, and SJK performed the analyses. RD contributed to the analysis of behavioral data. ABT, JYC and SPB wrote the manuscript with input from all authors.

---

**Figure S1.**
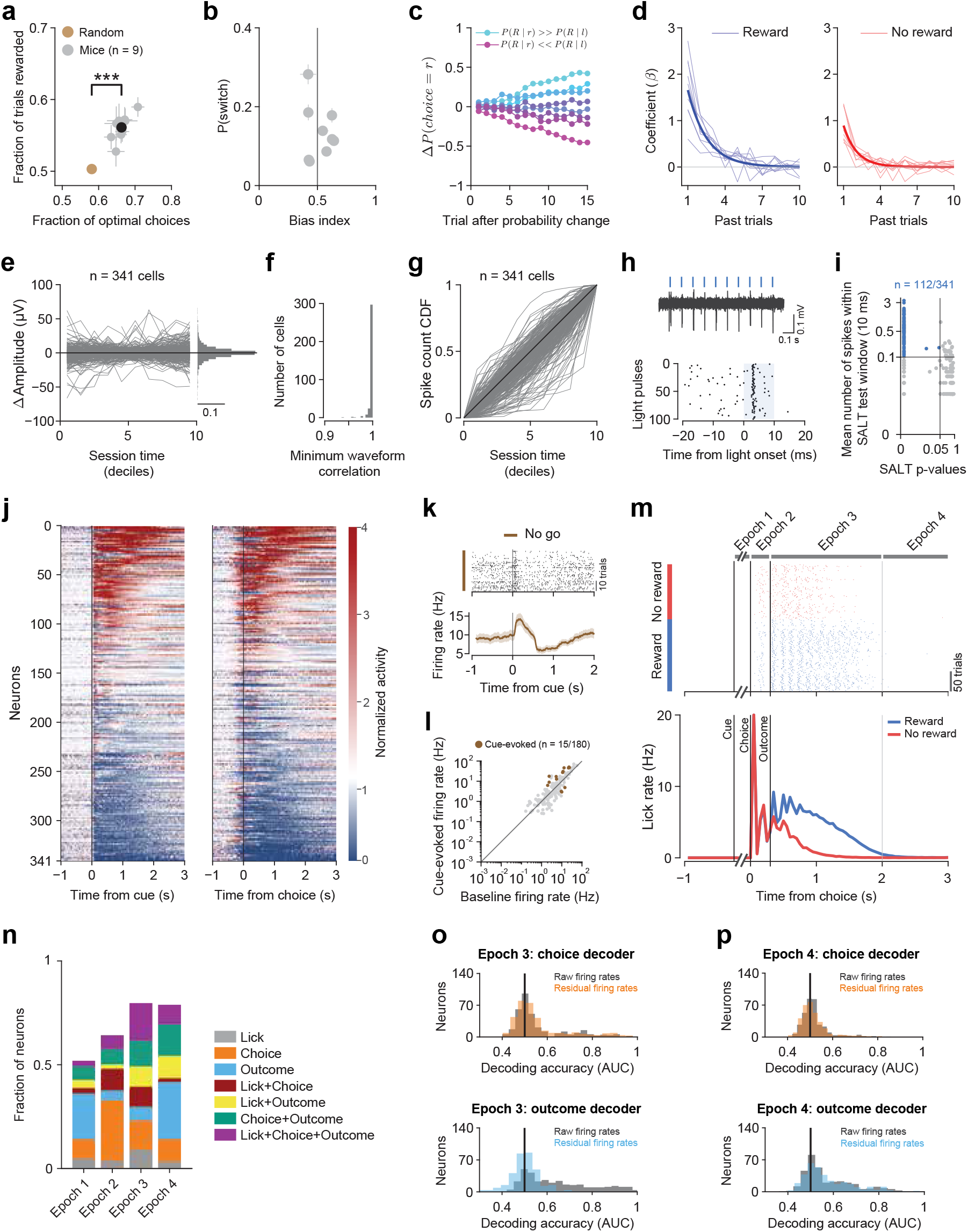
Behavioral performance and task-related activity of claustrum neurons. (**a**) Mice outperform a random-choice agent in both reward harvesting (Mice: 0.56 ± 0.018; Random: 0.50 ± 0.004; paired *t*-test = 11.25, *p* = 3.5 ×10^−6^) and optimal-choice fraction (Mice: 0.66 ± 0.021; Random: 0.58 ± 0.004; paired *t*-test = 12.15, *p* = 1.95 ×10^−6^), defined as the probability of selecting the higher-probability option (or either option when probabilities are equal). Gray points denote session averages for individual mice (*n* = 9); the black point shows the mean across mice; the orange point shows trial-matched random agent simulation. Some error bars (SEM) are obscured by symbols. (**b**) Switching probability as a function of individual choice bias. Each gray point shows the mouse-averaged switching probability (fraction of choice changes between consecutive trials) plotted against that mouse’s bias index (*N*_*r*_*/*(*N*_*r*_ + *N*_*l*_)) where *N*_*r*_ and *N*_*l*_ are the total number of rightward and leftward choices, respectively. A bias index of 0.5 indicates unbiased choice behavior, indicated by the vertical line. (**c**) Adaptation of choice probability following changes in reward contingencies. For each block transition, the probability of choosing right was computed over the first 15 trials after the switch. Curves show mean Δ*P* (*c*(*t*) = *r*). Blue traces correspond to blocks favoring the right option (*P* (*R* | *r*) *> P* (*R* | *l*)), and magenta traces to blocks favoring the left option (*P* (*R* | *l*) *> P* (*R* | *r*)). Mice systematically adjusted their choices in the appropriate direction, with faster and larger shifts observed with greater reward asymmetries. (**d**) Logistic regression coefficients predicting upcoming choice from past outcomes. Thin lines show coefficients fit separately for each mouse; bold curves show the exponential fit to population-averaged coefficients. (**e**) Mean spike amplitude for individual neurons (gray; *n* = 341) plotted across session time (deciles), showing minimal drift in waveform amplitude. The marginal histogram on the right shows the distribution of amplitudes across neurons (scale bar, probability density = 0.1). The black line denotes zero drift. (**f**) Histogram of the minimum waveform correlation between consecutive deciles for each neuron. All curated neurons exhibited high waveform similarity across time (minimum correlation *>* 0.9), indicating stable single-unit isolation. (**g**) Cumulative distribution functions (CDFs) of spike counts across session time (deciles) for individual neurons (*n* = 341), indicating unit presence throughout the session. The black line denotes the unity line corresponding to a uniform spike distribution across time. (**h**) Example optogenetic response of a claustrum neuron. Top, raw voltage trace showing light-evoked spiking following individual light pulses (blue ticks). Bottom, raster plot aligned to light onset, demonstrating reliable short-latency responses. (**i**) Mean number of spikes within a 10-ms window following light onset plotted against SALT *p*-values for all recorded neurons. Blue points indicate light-responsive neurons (*n* = 112*/*341 neurons); gray points indicate non-responsive neurons. The dashed line marks the significance threshold. (**j**) Heatmaps of mean activity, normalized to the 1 s pre-cue baseline, for all recorded claustrum neurons (*n* = 341 neurons, *N* = 9 mice) aligned to cue onset (left) or choice (right) for go trials. Neurons are sorted by their mean activity during the 3 s following cue onset (left) or choice (right). (**k**) Example neuron activity during no-go trials. Top, spike raster aligned to cue onset. Bottom, corresponding PSTH showing cue-evoked modulation in the absence of a behavioral response (shading, SEM). (**l**) Scatter plot of baseline versus cue-evoked firing rates across neurons recorded during 15 or more no-go trials without a licking response (*n* = 180*/*341 neurons). Brown points indicate significantly cue-responsive neurons (*n* = 15*/*180 neurons); gray points indicate non-responsive neurons. (**m**) Top, example choice-aligned lick raster from a single session, showing individual lick times for rewarded (blue) and unrewarded (red) trials. Vertical black lines mark: 1) auditory cue, 2) choice, 3) outcome. Task execution was divided into four behavioral epochs: Epoch 1: 0–100 ms following the cue; Epoch 2: 0–0.3 s following the choice, the period between choice and outcome delivery; Epoch 3: 0.3–2 s following the choice, the period following outcome in which *>* 99% of licks occur during task execution; and Epoch 4: 2–3 s following the choice, when *<* 1% of licks occur during task execution. Bottom, grand-average lick rate (*n* = 119 sessions, *N* = 9 mice; mean ± SEM across sessions) aligned to choice for rewarded (blue) and unrewarded (red) trials, computed in 50-ms bins. Mice licked significantly more during rewarded trials as compared to unrewarded trials during Epoch 3 (Rewarded: 9.7 ± 0.1 licks, Unrewarded: 4.1 ± 0.06 licks; paired *t*-test, *p* = 1.18 ×10^−83^). (**n**) Fraction of claustrum neurons (*n* = 341) significantly modulated by lick rate, choice, outcome and their combinations, in each behavioral epoch. (**o**) Top, histograms of single-neuron choice decoding accuracy (AUC) in Epoch 3 using a linear SVM applied to firing rates from individual claustrum neurons (*n* = 341). Gray histogram shows decoding based on raw firing rates. Colored histogram shows decoding based on residual firing rates, obtained after regressing out variance explained by lick rate and outcome. The vertical black line indicates chance performance (AUC = 0.5). Neurons with above-chance decoding were identified using shuffle-null testing (real AUC *>* 95% of shuffled AUCs; raw: *n* = 105*/*341 neurons; residual: *n* = 102*/*341 neurons; overlap: 77%), indicating that choice is significantly decoded from 30% of neurons during Epoch 3 after accounting for licking and outcome. Bottom, histograms of single-neuron outcome decoding accuracy (AUC) in Epoch 3 (raw: *n* = 214*/*341 neurons; residual: *n* = 57*/*341 neurons; overlap: 18%), indicating that outcome is significantly decoded from 17% of neurons during Epoch 3 after accounting for licking and choice. (**p**) Top, histograms of single-neuron choice decoding accuracy (AUC) in Epoch 4, after the overwhelming majority of licks have occurred (*>* 99% of licks during task execution), using a linear SVM shown as in (o) (raw: *n* = 55*/*341 neurons; residual: *n* = 56*/*341 neurons; overlap: 67%), indicating that choice is significantly decoded in 16% of neurons during Epoch 4 after accounting for lick rate and outcome. Bottom, histograms of single-neuron outcome decoding accuracy (AUC) in Epoch 4 (raw: *n* = 149*/*341 neurons; residual: *n* = 153*/*341 neurons; overlap: 89%), indicating that outcome is significantly decoded from 45% of neurons during Epoch 4 after accounting for lick rate and choice.

**Figure S2.**
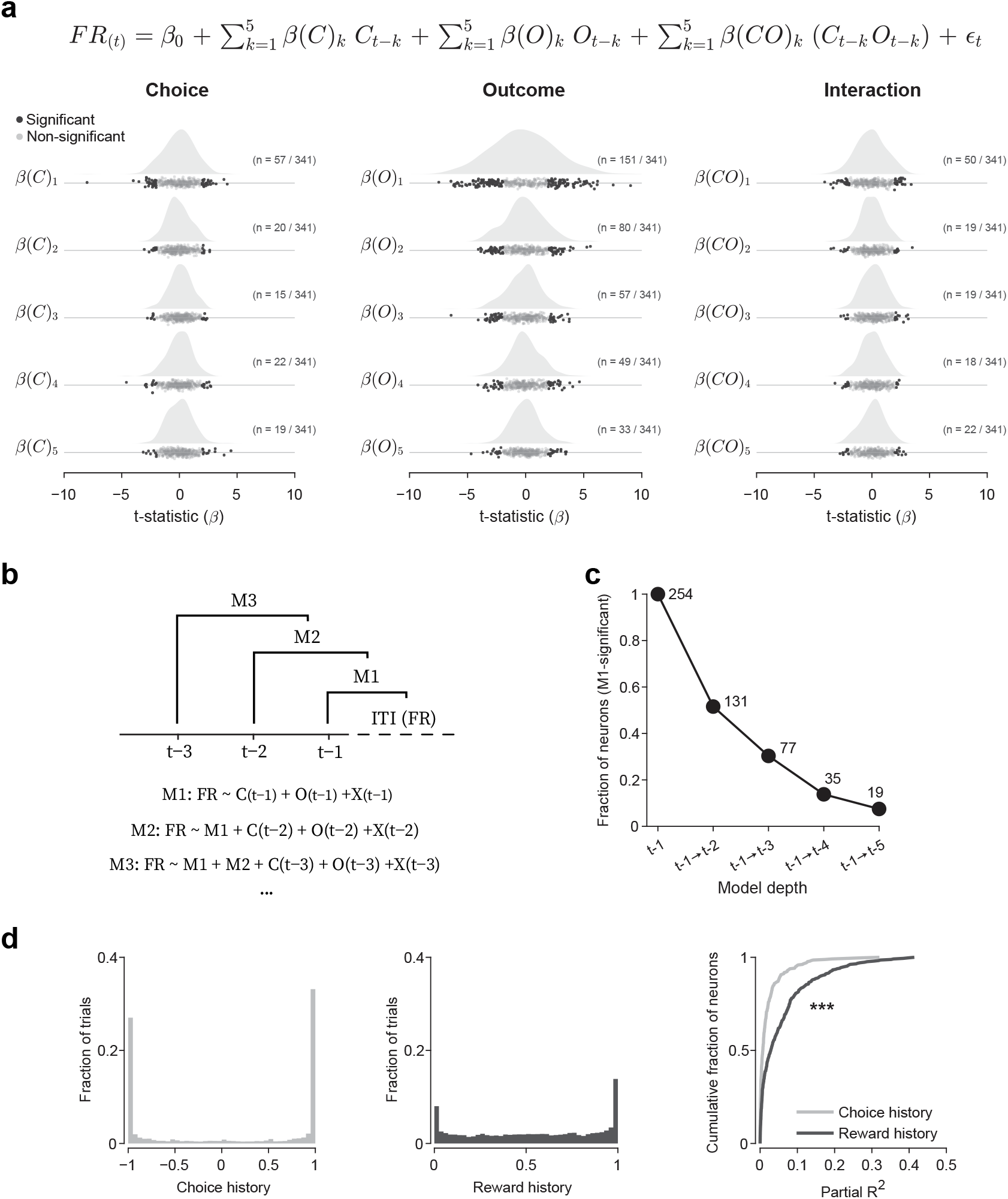
Lagged history representations of claustrum neurons. (**a**) Distribution of regression *t*-statistics for lagged choice (left), outcome (middle), and choice-by-outcome interaction terms (right) across lags from 1 to 5 trials back. For each lag, ridgeline plots show the density of *t*-values, overlaid with jittered points indicating individual neurons (*n* = 341). Dark points indicate neurons with significant coefficients (*α <* 0.05). Numbers denote the count of significant neurons out of the total recorded population at each lag. (**b**) Nested regression models sequentially incorporating terms from more trials in the past. Model M1 includes regressors for choice, outcome, and their interaction from the immediately preceding trial *t* − 1; M2 adds *t* − 2; M3 adds *t* − 3; and so on. (**c**) Fraction of neurons with significant Model M1 fits whose likelihood-ratio tests supported models incorporating progressively longer trial history. Values above each point indicate the number of neurons retained at each additional history lag from the population of 341 recorded neurons. (**d**) Distributions of choice-history (left) and reward-history (middle) values across behavioral sessions (*n* = 119 sessions, *N* = 9 mice). Cumulative distributions (right) of the unique variance explained (partial *R*^2^) by reward history versus choice history across neurons. Reward history accounted for significantly more unique variance than choice history (*n* = 341 neurons, Wilcoxon signed-rank test, *p <* 0.001).

**Figure S3.**
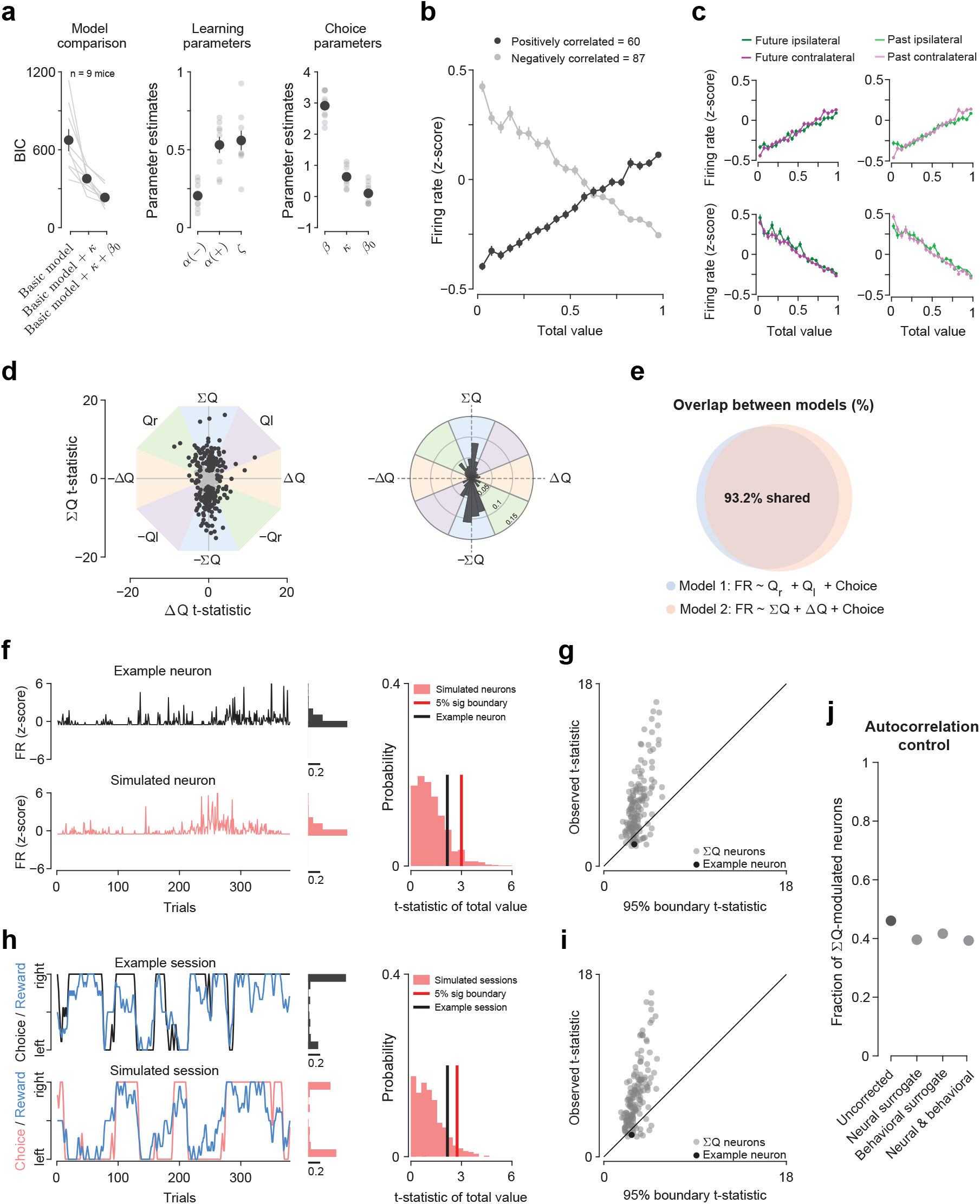
Total value representations of claustrum neurons. (**a**) Model comparison and parameter estimates. Left, Bayesian Information Criterion (BIC) values for three reinforcement-learning models of increasing complexity (Basic model; Basic + *κ*; Basic + *κ* + *β*_0_). Lines represent individual mice; points indicate group means ± SEM across mice. Middle, posterior mean estimates of learning parameters (*α*_(+)_, *α*_(−)_ and *ζ*) from the best-fitting model. Right, posterior mean estimates of choice parameters (*β, κ*, and *β*_0_). Gray points represent individual mice; black markers denote group means ± SEM. (**b**) Mean *z*-scored firing rates plotted as a function of Σ*Q* (bin size = 0.05) for Σ*Q* neurons with activity showing significant positive (dark gray; *n* = 60*/*147 neurons) or negative (light gray; *n* = 87*/*147 neurons) correlations with Σ*Q*. Error bars indicate ± SEM. (**c**) Choice invariance of Σ*Q* representations. Left, same analysis as in (b), split by upcoming choice (Ipsilateral, green; Contralateral, purple) for positively (top, *n* = 60*/*147 neurons) and negatively (bottom, *n* = 87*/*147 neurons) correlated Σ*Q* neurons. Σ*Q*-dependent activity did not differ significantly between upcoming choices (negatively correlated neurons sign-flipped for statistical testing; paired *t*-test, *t* = −1.5, *p* = 0.16). Right, same analysis as in (b), split by prior choice (Ipsilateral, light green; Contralateral, light purple), showing no significant difference in Σ*Q*-dependent activity between prior choices (negatively correlated neurons sign-flipped for statistical testing; paired *t*-test, *t* = −0.6, *p* = 0.56). (**d**) Axis-invariant regression analysis. Left, scatter plot of *t*-statistics for Σ*Q* versus Δ*Q* for all neurons (*n* = 341). Neurons with significant coefficients are shown in black (*p <* 0.05). Background shading denotes sectors corresponding to selective encoding axes. Right, polar histogram of preferred encoding angles, showing an over-representation along the Σ*Q* axis (*V* -test on folded angles, *p <* 0.001). (**e**) The majority of neurons identified as Σ*Q* neurons using the standard regression (Model 1: FR ∼ *Q*_*l*_ + *Q*_*r*_ +choice) were also classified as Σ*Q* neurons using the axis-invariant model (Model 2: FR ∼ Σ*Q* +Δ*Q* +choice). Overlap: n = 137*/*147 Model 1 Σ*Q* neurons (93.2%); n = 137*/*157 Model 2 Σ*Q* neurons (87.3%). (**f**) Neural surrogate control. Left, example neuron (top) and its matched amplitude-adjusted Fourier transform (AAFT) surrogate (bottom). The marginal histograms on the right of each trace show the distribution of firing rates across the session, demonstrating that the AAFT surrogate preserves the firing rate distribution of the example neuron (scale bar, probability density = 0.2). Right, distribution of surrogate Σ*Q t*-statistics (gray; 500 surrogates), with the 5% significance boundary indicated by the red vertical line. The example neuron’s Σ*Q t*-statistic (black vertical line) falls below this boundary, indicating that its apparent Σ*Q* correlation can be explained by intrinsic autocorrelation structure. (**g**) Comparison of each neuron’s observed Σ*Q t*-statistic versus its surrogate-derived 5% significance boundary for the population of Σ*Q* neurons defined using the axis-invariant model in (d) (Model 2: FR ∼ Σ*Q* + Δ*Q*+ choice). More than 85% of neurons lie above the unity line (*n* = 135*/*157 neurons), indicating significantly stronger Σ*Q* modulation than expected from intrinsic firing rate autocorrelations alone. (**h**) Behavioral surrogate control. Left, example behavioral session (top) and a matched surrogate simulation (bottom) preserving behavioral statistics while disrupting neural-behavioral relationships. The marginal histograms on the right of each session show the distribution of choices across the session, indicating differences in the overall session structure between the example and simulated sessions. Right, distribution of surrogate Σ*Q t*-statistics (gray; 500 surrogates), with the 5% significance boundary indicated by the red vertical line. The example neuron’s Σ*Q t*-statistic (black vertical line) falls below this boundary, indicating that its apparent Σ*Q* correlation can be explained by behavioral autocorrelation structure. (**i**) Comparison of each neuron’s observed Σ*Q t*- statistic versus behavioral surrogate 5% significance boundaries defined using the axis-invariant model in (d) (Model 2: FR ∼ Σ*Q* + Δ*Q*+ choice). Approximately 90% of neurons lie above the unity line (*n* = 142*/*157 neurons), indicating significantly stronger Σ*Q* modulation than expected from behavioral autocorrelation alone. (**j**) Fraction of Σ*Q* neurons from the population of recorded claustrum neurons retained after applying surrogate controls (Uncorrected: 46%, *n* = 157*/*341 neurons defined using Model 2; Neural surrogate correction: 40%, *n* = 135*/*341 neurons; Behavioral surrogate correction: 42%, *n* = 142*/*341 neurons; Neural and behavioral surrogate corrections: 39%, *n* = 134*/*341 neurons), indicating that approximately 40% of recorded neurons are Σ*Q* neurons after accounting for autocorrelation effects.

**Figure S4.**
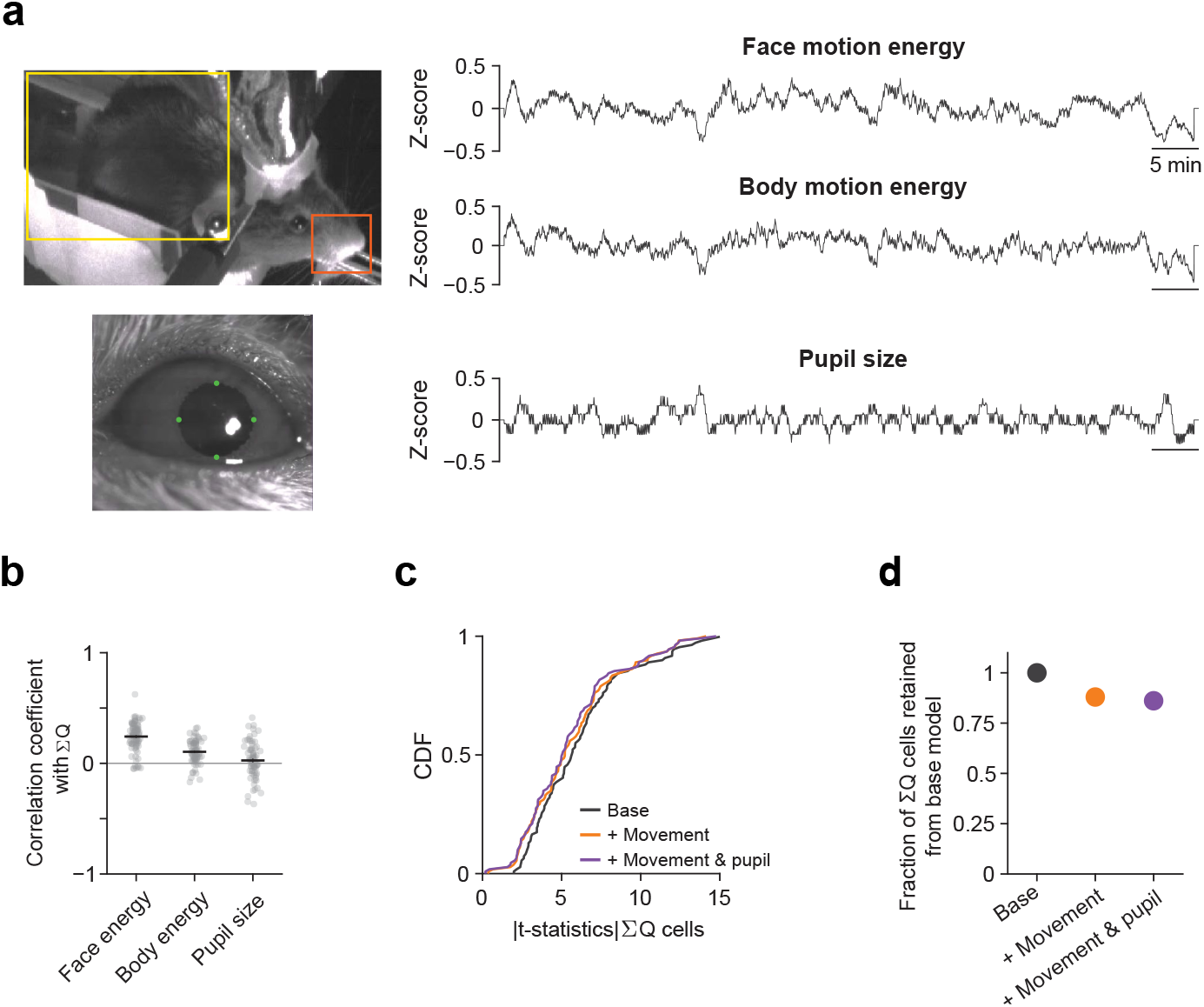
Movement and pupil dynamics do not account for total value representations in claustrum neurons. (**a**) Extraction of movement (top) and pupil (bottom) signals from synchronized behavioral videos. Left, example face region (orange), body region (yellow), and pupil tracking frame, as defined across all behavioral sessions with available videos (Face: *n* = 89*/*119 sessions; Body: *n* = 77*/*119 sessions; Pupil: *n* = 71*/*119 sessions). Right, *z*-scored traces of face motion energy, body motion energy, and pupil size over time from a sample behavioral session. (**b**) Correlation coefficient between each video-derived variable and Σ*Q* across sessions; horizontal lines show the mean ± SEM (Face energy: *r* = 0.24 ± 0.01, *n* = 89 sessions; Body energy: *r* = 0.10 ± 0.01, *n* = 77 sessions; Pupil size: *r* = 0.03 ± 0.02, *n* = 71 sessions). (**c**) Cumulative distributions of absolute Σ*Q t*-statistics across regression models of increasing complexity. The base model (black) is compared to models including face motion energy (+ Movement, orange) and including both movement and pupil size (+ Movement & Pupil, purple). Distributions are not significantly different from the base model (two-sample Kolmogorov–Smirnov (KS) tests: base model vs. base model + Movement, *p* = 0.61; base model vs. base model + Movement & Pupil, *p* = 0.4). (**d**) Fractions of Σ*Q* neurons identified by the base model that remain significant after including movement and pupil regressors. 87% of Σ*Q* neurons were retained after adding movement predictors (base + Movement: *n* = 117*/*134 Σ*Q* neurons among 271 recorded neurons with available movement videos). 85% of Σ*Q* neurons were retained after adding movement and pupil predictors (base + Movement & Pupil: *n* = 93*/*109 Σ*Q* neurons among 212 recorded neurons with available movement and pupil videos).

**Figure S5.**
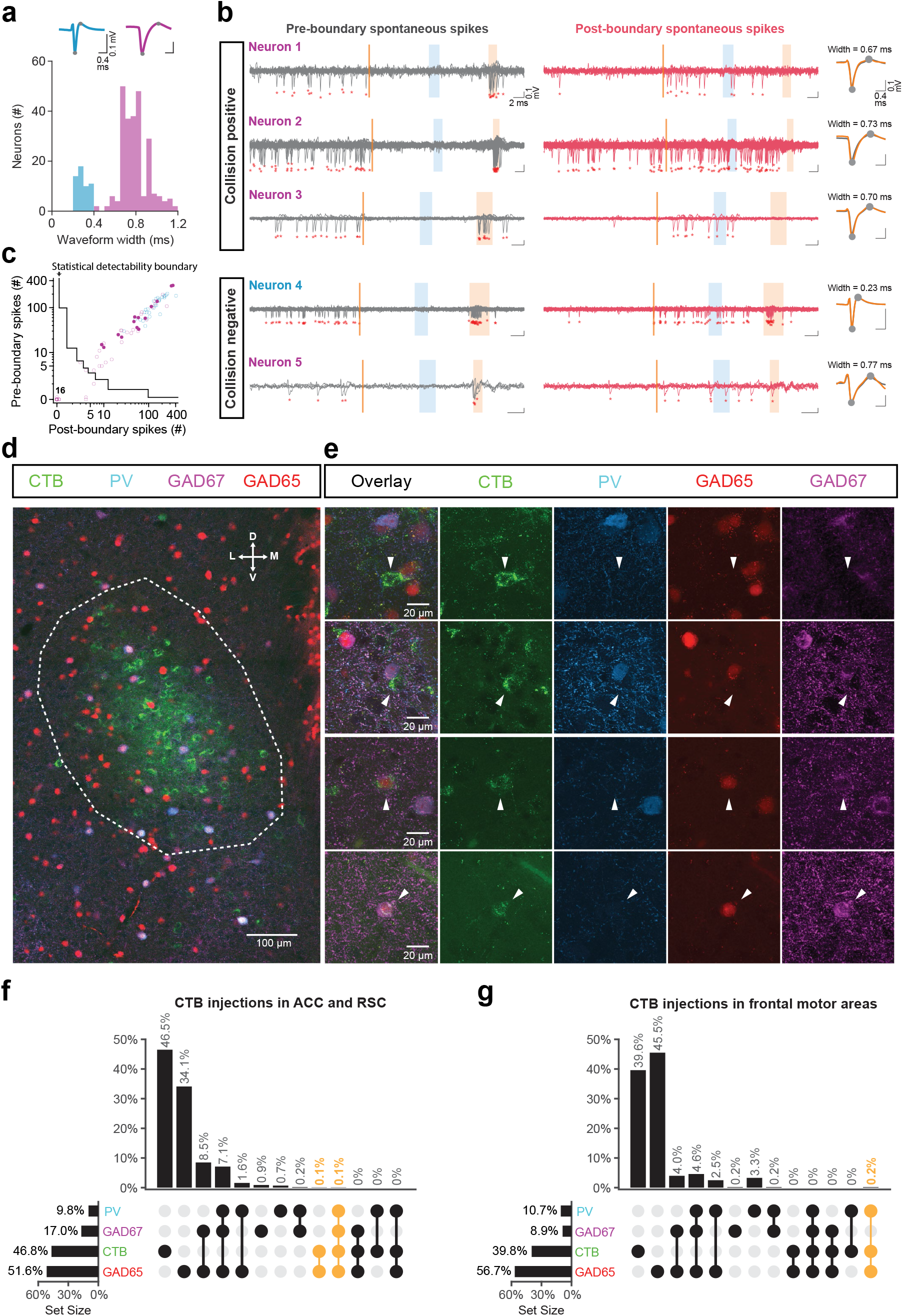
Claustrocortical neurons are wide-spiking neurons and excitatory. (**a**) Distribution of extracellular waveform widths of claustrum neurons recorded using a Neuropixels probe during collision test experiments (Narrow-spiking (NS): *n* = 53*/*300 neurons, 18%; Wide-spiking (WS): *n* = 247*/*300 neurons, 82%). Example NS and WS waveforms (top) illustrate the valley–peak measurements used to compute waveform width. (**b**) Representative neurons illustrating collision test outcomes. For each neuron, traces with spontaneous spikes occurring only in the pre-boundary window (left, gray) are compared with traces with spontaneous spikes occurring in the post-boundary window (right, red). For collision-positive neurons, the probability of detecting an evoked spike was significantly less when spontaneous spikes occurred after the collision boundary than before (neurons 1–3, WS neurons; Fisher’s exact test, *p <* 0.01). For collision-negative neurons, evoked spikes were reliably detected following spontaneous spikes occurring both before and after the collision boundary (neuron 4, NS neuron; Fisher’s exact test, *p* = 0.56), or the reduction in evoked spike probability that did not reach statistical significance (neuron 5, WS neuron; Fisher’s exact test, *p* = 0.048). Average spontaneous (gray) and evoked (orange) spike waveforms are shown to the right for each neuron, with waveform widths indicated. (**c**) Theoretical detectability analysis for collision testing based on spike counts alone. Each point represents a recorded neuron plotted by the number of spontaneous spikes occurring before the collision boundary (pre-boundary) and after the boundary (post-boundary). The black stepwise curve denotes the statistical detectability boundary, derived under idealized assumptions that photostimulation always evokes an action potential following pre-boundary spikes while post-boundary spontaneous spikes always abolish evoked spikes. For each combination of pre- and post-boundary spike counts, a Fisher’s exact test (*α* = 0.01) was used to determine whether collision detection was statistically powered. Neurons falling below the boundary are underpowered for collision detection (WS: *n* = 21*/*59 neurons, NS: *n* = 0*/*29 neurons). (**d**) Low-magnification image of a coronal section of the claustrum showing neurons retrogradely labeled from ACC and RSC (green), immunostained for parvalbumin (PV; blue) and the inhibitory neuron marker GAD67 (magenta), and expressing GAD65-mCherry (red). Scale bar: 100 *µ*m. (**e**) Higher-magnification images of the claustrum highlighting four example retrogradely labeled neurons, showing collocalization of markers. Scale bar: 20 *µ*m. (**f**) UpSet plot illustrating the intersectional analysis of neurons retrogradely labeled from ACC and RSC, immunopositive for PV and GAD67, and GAD65-mCherry-positive (red). Eight of the 3501 CTB-positive claustrocortical neurons were annotated as PV-positive; 17 of the 3501 CTB-positive neurons were annotated as GAD67-positive and/or GAD65-mCherry-positive (*N* = 4 mice). (**g**) Same as (f), but for claustrocortical neurons retrogradely labeled from frontal motor areas. One of the 227 retrogradely labeled claustrocortical neurons was annotated as PV-positive; 1 of the 227 CTB-positive neurons was annotated as GAD65-mCherry-positive (*N* = 4 mice).

**Figure S6.**
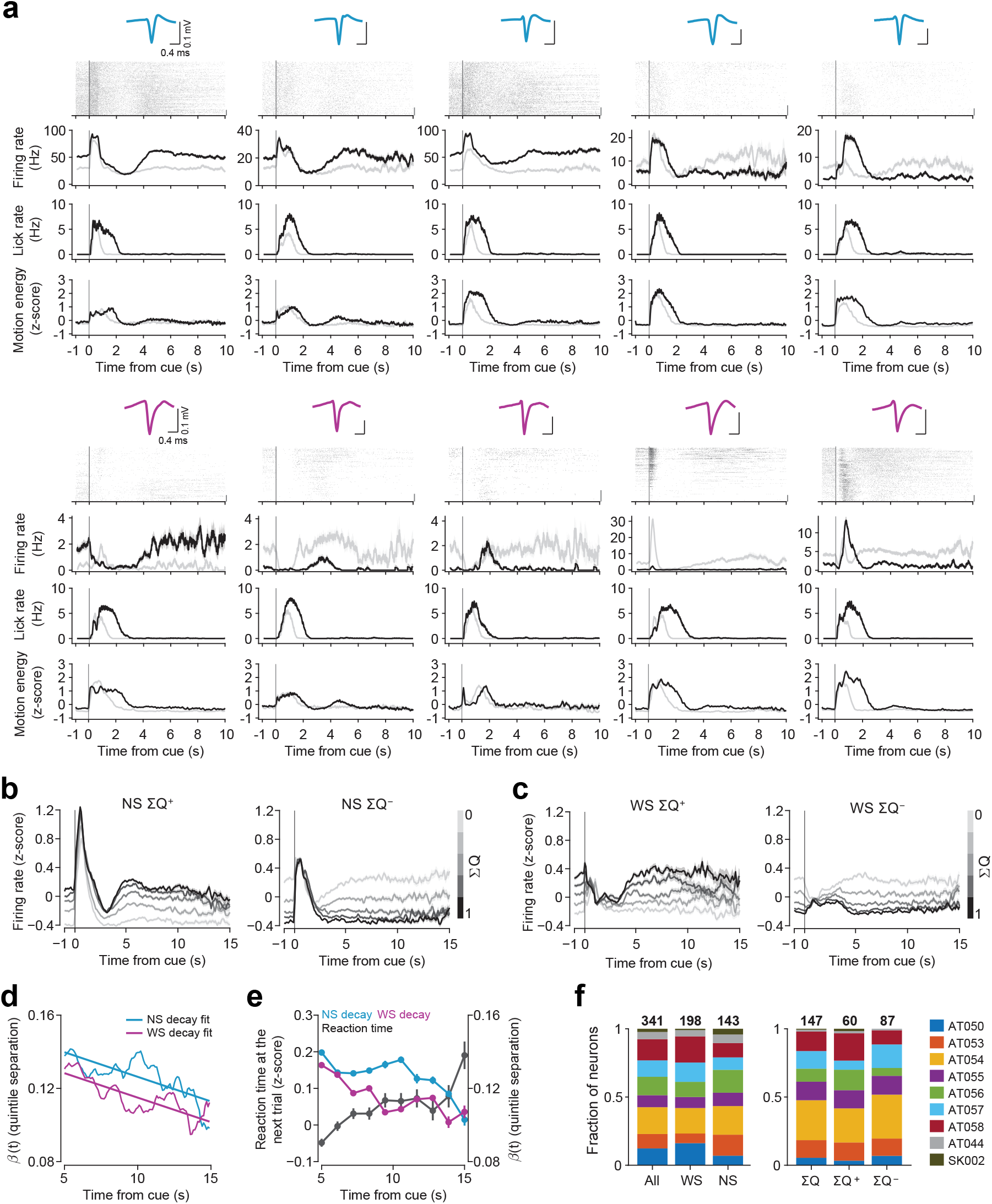
Sustained total value signals in narrow-spiking (NS) and wide-spiking (WS) claustrum neurons. (**a**) Cue-aligned raster plots with trials sorted by increasing Σ*Q*, followed by PSTHs, lick rate and face motion energy for low Σ*Q* (light) and high Σ*Q* (dark) trials (median split) for five example NS neurons (top) and five example WS neurons (bottom). Each neuron’s average spike waveform is shown above the plots. (**b**) Population PSTHs for NS neurons with activity significantly positively (left) and negatively (right) correlated with Σ*Q*. Z-scored firing rates are split across Σ*Q* quintiles (light to dark); shading indicates SEM. (**c**) As in (b), but for WS neurons. (**d**) *β*(*t*) (quintile separation) plotted versus ITI length for both NS (blue) and WS (magenta) neurons. Solid lines indicate linear fits, showing comparable decay rates (linear model, Time ×Cell type interaction: *t* = −0.01, *p* = 0.99). (**e**) Reaction time on the upcoming trial (gray, left axis), NSΣ*Q* stability (*β*(*t*); blue, right axis) and WS Σ*Q* stability (*β*(*t*); magenta, right axis) plotted against the ITI length. *β*(*t*) and reaction time are negatively correlated for both cell types (NS: *r* = −0.83; WS: *r* = −0.79). (**f**) Left, fraction of all neurons, and neurons classified as NS and WS, recorded from each mouse. Right, fraction of all Σ*Q* neurons, and neurons positively and negatively correlated with Σ*Q*, recorded from each mouse. Colors represent individual mice.

